# 27-hydroxylation of oncosterone by CYP27A1 switchs its activity from pro-tumor to anti-tumor

**DOI:** 10.1101/2023.10.10.560948

**Authors:** Silia Ayadi, Silvia Friedrichs, Regis Soulès, Laly Pucheu, Dieter Lütjohann, Sandrine Silvente-Poirot, Marc Poirot, Philippe de Medina

## Abstract

Oncosterone (6-oxo-cholestane-3β,5α-diol; OCDO) is an oncometabolite and a tumor promoter on estrogen receptor alpha positive breast cancer (ER(+) BC) and triple negative breast cancers (TN BC). OCDO is an oxysterol formed in three steps from cholesterol: 1) oxygen addition at the double bond to give α- or β-isomers of 5,6-epoxycholestanols (5,6-EC), 2) hydrolyses of the epoxide ring of 5,6-ECs to give cholestane-3β,5α,6β-triol (CT), and 3) oxidation of the C6 hydroxyl of CT to give OCDO. On the other hand, cholesterol can be hydroxylated by CYP27A1 at the ultimate methyl carbon of its side chain to give 27-hydroxycholesterol (27HC), which is a tumor promoter for ER(+) BC. It is currently unknown whether OCDO and its precursors can be hydroxylated at position C27 by CYP27A1, as is the impact of such modification on the proliferation of ER(+) and TN BC cells. We investigated, herein, whether 27-hydroxylated-5,6-ECs, -CT and -OCDO exist as metabolites and can be produced by cells expressing CYP27A1. We report, for the first time, that these compounds exist as metabolites in human. We give pharmacological and genetic evidences that CYP27A1 is responsible for their production. Importantly, we found that 27-hydroxy-OCDO (27H-OCDO) inhibits BC cells proliferation and blocks OCDO and 27-HC induced proliferation in BC cells, showing that this metabolic conversion commutes the proliferative properties of OCDO into antiproliferative ones. These data suggest an unprecedented role of CYP27A1 in the control of breast carcinogenesis by inhibiting the tumor promoter activities of oncosterone and 27-HC.

## INTRODUCTION

Oxysterols constitute a growing family of oxygenated cholesterol metabolites that impact on a plethora of fatal diseases including atherosclerosis, cancer, neurodegenerative diseases and aging (1–4). Depending on their chemical structures, these metabolites exert distinct biological properties through various molecular mechanisms that could involve either nuclear receptors (e.g., Liver-X-Receptor, LXR; retinoid-related orphan receptor, ROR; estrogen receptor alpha, ERα; glucocorticoid receptor alpha, GRα) (5–8), G-protein coupled receptors (e.g., smoothened, SMO; CXC-motif-chemokine receptor 2, CXCR2; Epstein-Barr virus-induced gene 2, EBI2) (9–13), enzymes (e.g., sterol O-acyltransferase/Acyl-coA cholesterol acyltransferase, SOAT/ACAT; cholesterol-5,6-epoxide hydrolase, ChEH; and 3-hydroxy-3-methylglutaryl-coenzyme A reductase, HMG-CoAR; 11β-hydroxysteroid dehydrogenase type 2, (HSD2) (14–16) or transporters (e.g., ions transporters, insulin-induced gene, INSIG; sterol or oxysterol binding proteins (ORP) proteins; Nieman-Pick C1 and C1 like 1, (NPC1 and NPC1L1) (17, 18). Several studies highlight that the deregulation of the metabolism of oxysterols plays a pivotal role in the development of cancers and more particularly breast cancers (BC) (1, 19–21). Indeed, two tumor promoters and one tumor suppressor were identified. 27-hydroxycholesterol (27HC) (Fig. 1A) is produced by CYP27A1 and has been shown to be a tumor promoter in estrogen receptor alpha positive BC (ER(+) BC). It is one of the major circulating side-chain oxysterol (16, 22, 23), but while its tumor promoter activity was established in mice (22, 23) and inverse association was found between high circulating 27HC level and breast cancer risk (19, 24–26). 6-oxo-cholestan-3β,5α-diol (OCDO (Fig. 1B), denoted oncosterone), is a B-ring oxysterol oncometabolite identified in BC and was reported to be a tumor promoter for ER(+) BC and triple negative BC (TN-BC) (5, 27, 28). Dendrogenin A (Fig. 1B) a B-ring oxysterol conjugate produced by healthy tissues including breast that was reported to display tumor suppressor properties(3, 27, 29–34). Oncosterone is a tertiary metabolite of cholesterol (Fig. 1B). Cholesterol is first epoxidized at the level of its delta-5,6 double bond to give 5α,6α-epoxycholestanol (5,6α-EC) and 5β,6β-epoxycholestanol (5,6β-EC) (Fig. 1C) (35). This addition reaction can be done through lipid peroxidation or via cytochrome p450 (35). In cancer cells, 5,6-ECs are not metabolized into DDA but are mainly metabolized into cholestane-3β,5α,6β-triol (CT) by the ChEH (36, 37), which is next metabolized into OCDO (Oncosterone) by HSD2 (5, 28) (Fig. 1C). The expression levels of OCDO-forming enzymes (ChEH and HSD2) are upregulated in BC compared to normal adjacent tissues and their expression negatively correlated with patient survival (5, 38). The B-ring oxysterol 7-ketocholesterol was reported to be hydroxylated at position C27 by CYP27A1 to give 27H-7ketocholesterol (39) (Fig. 1D) suggesting that other B-ring oxysterols could be 27-hydroxylated by this enzyme. In the present study, we evaluated whether OCDO and its precursors can undergo a 27-hydroxylation in cancer cells expressing CYP27A1 and compares the proliferative properties of 27H-OCDO on ER(+) and TN BC cells with OCDO and 27HC. We show that OCDO, CT and 5,6-ECs can be 27-hydroxylated by CYP27A1 and reveals a metabolic switching of the pro-tumor activity of OCDO to an anti-tumor activity in BC cells expressing CYP27A1 through the hydroxylation of OCDO at position C27.

**Figure 1:**
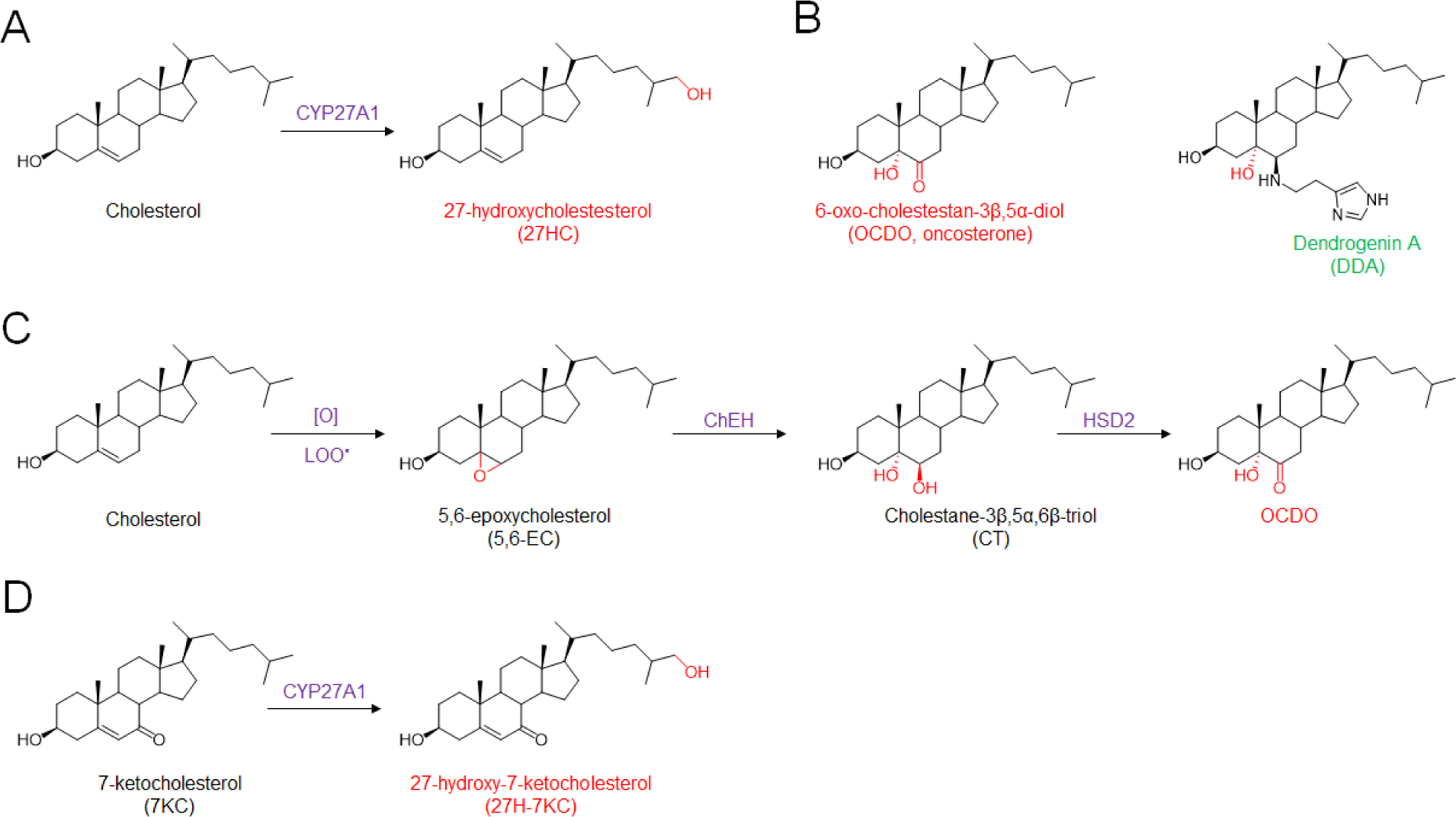
A) chemical structure of cholesterol metabolites with tumor promoter properties in breast cancers (red) and tumor suppressor properties (green). Metabolic pathways leading to the production of OCDO (B) and 27HC (C). [O] and LOO∙ represent respectively oxidative stress and lipid hydroperoxides radicals involved in the epoxidation of delta-5,6 double bond of cholesterol

## MATERIALS and METHODS

### Chemicals

[4-^14^C]-cholesterol (52 mCi/mmol) was purchased from Perkin Elmer (Whaltham, USA). [4-^14^C]-5,6α-epoxycholestan-3β-ol, [4-^14^C]-5,6β-epoxycholestan-3β-ol, [4-^14^C]-cholestane-3β,5α,6β-triol and [4-^14^C]-6-oxo-cholestan-3β,5α-diol were synthetized as previously described (15, 40, 41). (25R)-cholest-5-ene-3β,26-diol (27-hydroxycholesterol; 27HC), cholest-5-ene-3β,25-diol (25-hydroxycholesterol; 25HC), 6-oxo-cholestan-3β,5α-diol (OCDO; oncosterone) was from Steraloids (Newport, US) and bicalutamide and tamoxifen were from Sigma-Aldrich (Merck KGaA, Darmstadt, Germany). Other chemicals were from Sigma-Aldrich (Merck KGaA, Darmstadt, Germany). The advancement of the reactions was monitored by thin layer chromatography (TLC) in conditions were reactants and products are separated on Merck silica gel 60 F254 (0.040-0.063 mm) plates. Reactants and products were revealed after spraying the plates with sulfuric acid/methanol (1:1) and heating. The purification of newly synthesized compounds was performed by flash chromatography on a CombiFlash NextGen 300 (Serlabo, Entraigues sur la sorgues, France). ^1^H NMR and ^13^C NMR spectra were recorded on Bruker spectrometers (Avance III HD 400 MHz, 500 MHz or NEO 600 MHz, (Bruker BioSpin GmBH; Rheinstetten, Germany). Solutions of new compounds were prepared in deuterochloroform (CDCl_3_), deuterated acetone (acetone d_6_) or deuterated methanol (CD_3_OD). Deuterated solvents were from from Euriso-top (St-Alban, France) and were used as internal references. Mass determination of new compounds were recorded in positive mode on a DSQ II (Thermo Fisher Scientific, Les Ullis, France) using chemical ionization (NH_3_ as ionization gas). All the organic layers were dried using anhydrous MgSO_4_.

### Chemical syntheses

The chemical synthesis of the new compounds is summarized on Fig. 2.

**Figure 2:**
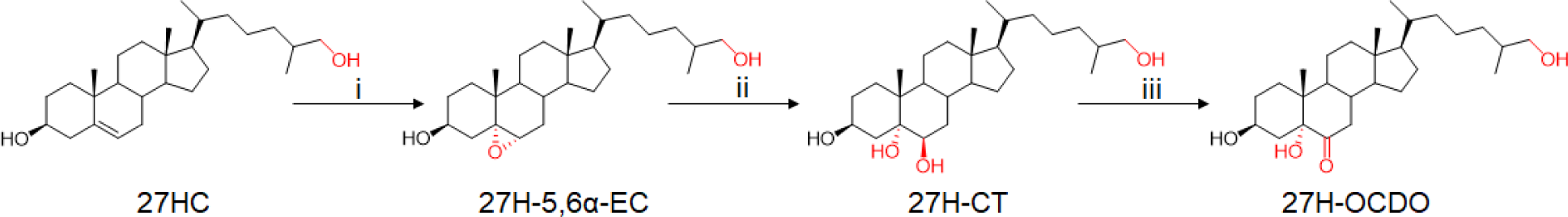
Synthesis of 27H-5,6α-EC, -CT and -OCDO. i : 1.5 equiv mCPBA, CHCl3, rt, 3h (90%); ii : HClO_4_, THF/H_2_O/acetone, rt, 30 min (38%); iii : 25 equiv NBS, diethylether/MeOH/H_2_O, rt, 3h (65%).

All compounds have been registered at LIPID MAPS® (https://www.lipidmaps.org/). Their ID is given as supplementary data S1.

#### Synthesis of 27-hydroxy-5,6-epoxycholestan-3β-ol (27H-5,6-EC)

To a solution of 27HC (40 mg; 0.1 mmol) in CHCl_3_ (2 ml) a solution of meta-chloroperbenzoic acid (0.15 mmol; 1.5 equiv) in CHCl_3_ (1 ml) was added. The mixture was stirred at room temperature (rt) for 2 h, diluted with 15 ml of chloroform and washed sequentially with Na_2_SO_4_ (10% aqueous solution [aq. sol.]), NaHCO_3_ (5% aq. sol.) and brine (water saturated with NaCl). The organic layer was dried over MgSO_4_, filtered and evaporated. The white crude product was purified by flash chromatography (CombiFlash NextGen) on a silica column (4G; column volume (CV) = 4.8 ml) using an ethyl acetate gradient in chloroform (0% EtOAc for 2 CV, then to 100% EtOAc in 15 CV and 100% EtOAc for 5 CV; flow rate = 13 ml/min) with an elution at 15-16 CV. The mixture of 27H-5,6α-EC (83%) and 27H-5,6β-EC (17%) were obtained as an amorphous white solid (90% yield). Rf (EtOAc) = 0.47. ^1^H-NMR (500 MHz, CDCl_3_): δ = 3.87-3.90 (1H, m, H-3, α-isomer), 3.66-3.72 (1H, m, H-3, β-isomer), 3.48-3.51 (1H, m, H-27, isomers α+β), 3.39-3.43 (1H, m, H-27, isomers α+β), 3.05 (1H, d, J= 2.3 Hz, H-6, β-isomer), 2.9 (1H, d, J=4.4, H-6, α-isomer), 1.05 (3H, s, H-19, α+β-isomers), 0.88-0.91 (6H, d, H-26 and H-21, α and β isomers), 0.63 (3H, s, H-18, β-isomer), 0.6 (3H, s, H-19, α-isomer). ^13^C-NMR (500 MHz, CDCl_3_): 69.58 (C-3, β-isomer), 68.89 (C-3, α-isomer), 68.67 (C-27, α+β isomers), 65.82 (C-5, α-isomer), 63.85 (C-6, β-isomer), 63.07 (C-5, β-isomer), 59.43 (C-6, α-isomer), 56.99, 56.37, 56.33, 55.99, 51.47, 42.7, 42.49, 42.44, 42.38, 40.02, 39.56, 37.38, 36.24, 35.93, 35.82, 35.00, 33.65, 32.76, 32.54, 30.04, 29.92, 28.96, 28.331, 28.24, 24.32, 24.18, 23.55, 22.14, 20.79, 18.77, 18.74, 17.19, 16.63, 16.07, 12.01, 11.91. MS (DCI-NH_3_) m/z: 436.4 [M-NH_4_^+^]; 419.4 [MH^+^]; 401.4 [MH^+^-H_2_O]; 383.4 [MH^+^-2H_2_O].

#### Synthesis of 27-hydroxy-cholestane-3β,5α,6β-triol (27H-CT)

To a solution of 27H-5,6-EC (87% α isomer; 13% β isomer determined by ^1^H-NMR) (20 mg; 0.048 mmol) in 3.25 ml of tetrahydrofurane/water/acetone (v/v/v; 2:1:0.25) was added perchloric acid (0.25 ml). The mixture was stirred at room temperature for 30 minutes diluted with 15 ml of butanol and washed sequentially with NaHCO_3_ (5% aq. sol.) and brine. The organic layer was dried over anhydrous MgSO_4_, filtered and evaporated. The white crude product was solubilized in a minimum of methanol. 27H-CT was purified by flash chromatography on a silica column (12G column volume (CV)=16.8 ml) using an acetone gradient in chloroform (0% acetone for 2 CV, then to 100% acetone in 15 CV and 100% acetone for 5 CV; flow rate= 30 ml/min) with an elution at 12-15 CV. 27H-CT was obtained as an amorphous white solid (38% yield). Rf (EtOAc)=0.1. ^1^H NMR (500 MHz, acetone d_6_/ethanol d_6_) δH = 4.00 (m, 1H, H-3), 3.48 (Broad t, 1H, H-6), 3.36-3.39 (m, 1H, H-27), 3.27-3.30 (m, 1H, H-27), 1.5 (s, 3H, H-19), 0.92 (d, 3H, J=6.6 Hz, H-21), 0.86 (d, 3H, J=6.7 Hz, H-26), 0.69 (s, 3H, H-18). ^13^C-NMR (400 MHz, acetone-d_6_/ethanol-d_6_): 75.12 (C-6), 74.68 (C-5), 67.05 (C-27), 66.69 (C3), 56.34, 45.17, 42.61, 40.72, 40.22, 38.11, 36.11, 35.83, 34.50, 33.54, 32.27, 30.90, 30.34, 28.07, 24.01, 23.29, 21.02, 18.15, 16.11, 11.58. MS (DCI-NH_3_) m/z: 454.3 [M-NH_4_^+^]; 436.3 [M-NH_4_ ^+^-H_2_O]; 418.3 [M-NH_4_^+^-2H_2_O]; 401.4 [MH^+^-2H_2_O].

#### Synthesis of 27-hydroxy-6-oxo-cholestan-3β,5α-diol (27H-OCDO)

27H-CT (20 mg; 0.046 mmol) was dissolved in diethylether (4.5 ml), methanol (0.75 ml) and water (0.75 ml). N-bromosuccinimide (1.14 mmol; 25 eq) was added and the reaction mixture was stirred at room temperature for 2 h. Diethylether (15 ml) was added and the organic layer was washed with brine. Then, the aqueous layer was washed with CHCl_3_. Organic layers were pooled, dried under MgSO_4_ and evaporated. The crude product was purified by flash chromatography on a silica column (4G; column volume (CV)=4.8 ml) using an ethyl acetate gradient in chloroform (0% EtOAc for 3 CV, then to 100% EtOAc in 20 CV and 100% EtOAc for 5 CV; flow rate= 13 ml/min) with an elution at 20-21 CV. 27H-OCDO was obtained as an amorphous white solid (65% yield). Rf (EtOAc)= 0.39. 1H-NMR (500 MHz, acetone-d_6_): δ= 3.92-3.96 (1H, m, H-3), 3.32-3.46 (2H, m, H-27), 2.78 (1H, t, J=12.5 Hz, H-7), 0.96 (3H, d, J= 6.6 Hz, H-21), 0.89 (3H, d, J=6.7 Hz, H-26), 0.8 (3H, s, H-19), 0.71 (3H, s, H-18) ppm. ^13^C-NMR (400 MHz, acetone d6): 212.24 (C-6), 80.94 (C-5), 68.20 (C-27), 67.06 (C-3), 57.49, 57.17, 45.40, 44.01, 43.18, 42.49, 40.82, 38.23, 37.17, 37.09, 36.86, 36.62, 34.58, 31.65, 30.84, 28.94, 24.71, 24.32, 22.31, 19.16, 17.20, 14.41, 12.51. MS (DCI-NH_3_) m/z : 452.4 [M - NH_4_^+^]; 434.4 [M-NH_4_^+^-H_2_O].

#### Synthesis of 25-hydroxy-5,6-epoxycholestan-3β-ol (25H-5,6-EC)

25H-5,6-EC was produced as described for the chemical synthesis of 27H-5,6-EC starting from commercially available 25HC. 25H-5,6-EC was obtained as an amorphous white solid (90% yield). Rf (EtOAc)=0.47. ^1^H-NMR (600 MHz, CDCl_3_): δ= 3.87-3.93 (1H, m, H-3, α-isomer), 3.66-3.72 (1H, m, H-3, β-isomer), 3.05 (1H, d, J= 2.3 Hz, H-6, β-isomer), 2.9 (1H, d, J=4.4, H-6, α-isomer), 1.06 (3H, s, H-19, α+β-isomers), 0.9-0.91 (3H, d, J= 6.6 Hz, H-21, α and β isomers), 0.64 (3H, s, H-18, β-isomer), 0.61 (3H, s, H-18, α-isomer). ^13^C-NMR (600 MHz, CDCl_3_): 71.12 (C-25), 69.46 (C-3, β-isomer), 68.76 (C-3, α-isomer), 65.68 (C-5, α-isomer), 63.71 (C-6, β-isomer), 62.93 (C-5, β-isomer), 59.29 (C-6, α-isomer), 56.85, 56.23, 56.13, 55.81, 51.13, 44.4, 42.56, 42.37, 42.32, 42.25, 39.88, 39,84, 39.42 37.24, 36.41, 35.73, 35,69, 34.87, 32.62, 32.41, 31.12, 31.07, 29.91, 29,78, 29.39, 29.33, 29.23, 29.19, 28.83, 28.17, 28.10, 24.19, 24.05, 22.00, 20.81, 20.74, 20.65, 18.65, 18.62, 17.06, 15.94, 11.87, 11.77. MS (DCI-NH_3_) m/z: 436.4 [M-NH_4_^+^]; 419.4 [MH^+^]; 401.4 [MH^+^-H_2_O]; 383.4 [MH^+^-2H_2_O].

#### Synthesis of 25-hydroxy-cholestane-3β,5α,6β-triol (25H-CT)

25H-CT was produced as described above for the chemical synthesis of 27H-CT starting from 25H-5,6-EC. The reaction mixture was maintened at 0°C rather than room temperature. 25H-CT was obtained as an amorphous white solid (25% yield). Rf (EtOAc)= 0.1. ^1^H-NMR (CD_3_OD) δH (ppm) = 4.01 (m, 1H, H-3), 3.45 (t, 1H, H-6), 1.17 (s, 6H, H-26 and H-27), 1.16 (s, 3H, H-19), 0.95 (d, 3H, J = 6.5 Hz, H-21), 0.71 (s, 3H, H-18). ^13^C-NMR (CD_3_OD) δC = 76.76 (C-5), 76.45 (C-6), 71.45 (C-25), 68.27 (C3), 57.59, 57.38, 46.50, 45.26, 43.87, 41.42, 41.35, 39.24, 37.07, 35.20, 33.42, 31.60, 31.54, 29.29, 29.24, 29.10, 25.16, 22.25, 21.82, 19.20, 17.29, 12.60. MS (DCI-NH_3_) m/z : 454.3 [M - NH_4_^+^]; 436.3 [M-NH_4_^+^-H_2_O]; 418.3 [M-NH_4_ ^+^-2H_2_O]; 401.4 [MH^+^-2H_2_0].

#### Synthesis of 25-hydroxy-6-oxo-cholestan-3β,5α-diol (25H-OCDO)

25H-OCDO was produced as described above for the chemical synthesis of 27H-OCDO starting from 25H-CT to give the expected product as a white amorphous solid (65% yield). Rf (EtOAc)=0.39. ^1^H-NMR (500 MHz, acetone d6): δ= 3.89-3.96 (1H, m, H-3), 2.76 (1H, t, J=12.5 Hz, H-7), 1.14 (6H, s, H26 and H-27), 0.96 (3H, d, J= 6.6 Hz, H-21), 0.78 (3H, s, H-19), 0.7 (3H, s, H-18) ppm. ^13^C-NMR (500 MHz, acetone d6) : 212.24 (C-6), 80.95 (C-5), 70.24 (C-25), 67.08 (C-3), 57.51, 57.21, 45.45, 45.43, 44.04, 43.20, 42.52, 40.86, 38.25, 37.58, 37.22, 36.75, 31.69, 30.86, 28.96, 24.73, 22.33, 21.64, 19.20, 17.20, 14.42, 12.52. MS (DCI-NH_3_) m/z: 435.4 [MH+]; 452.4 [M-NH_4_+].

### Cell culture

MCF7 (human ER(+) BC), MDA-MB-231 (human TN BC, MDA-231), MDA-MB-468 (human TN BC, MDA-468) and HepG2 (human hepatoblastoma) cells were from the American Type Culture Collection (ATCC) and cultured until passage 30, HepG2 cells until passage 50. HEK293FT (R7007) were from Thermo Fisher Scientific (Waltham, MA, USA) and were maintained in DMEM (Thermo Fisher Scientific, Waltham, MA, USA) supplemented with 10% FBS, 0.1 mM MEM non-essential amino acids, 2 mM glutamine, 1mM sodium pyruvate, 1% penicillin/streptomycin, and 500 μg/ml geneticin. Cell lines were tested once a month for mycoplasma contamination using Mycoalert® Detection Kit (Lonza, Basel, Switzerland). MCF7 and MDA-468 cells were grown in RPMI 1640 medium (Thermo Fisher Scientific, Waltham, MA, USA) supplemented with 5 % and 10% FBS (Dutscher, Bernolsheim, France) respectively. MDA-231 and HepG2 cells were grown in DMEM and EMEM respectively (Thermo Fisher Scientific, Waltham, MA, USA) supplemented with 10 % FBS. For 27HC treatments cells were grown in a medium without phenol red. All cell media were supplemented with penicillin and streptomycine (50 U/ml each). Cell lines were cultured in a humidified atmosphere with 5% CO_2_ at 37 °C.

### Side-chain cholesterol hydroxylase expression analysis

CYP27A1 gene expression analysis for different tissues from patients were downloaded from the Genotype-Tissue Expression (GTEx) database (GTEx Portal).

CYP27A1, CH25H and CYP46A1 expression on paired normal adjacent tissues, breast cancer and metastasis, data were retrieved from the TNM plot database ([TNMplot](https://tnmplot.com/analysis/)). The human protein Atlas database was used for the analysis of CYP27A1, CH25H and CYP46A1 expression in cancer cell lines (https://www.proteinatlas.org/).

### Prognosis value of HSD2 and CYP27A1 expression on breast cancer

Correlation between breast cancer patient survival and the mRNA expression of HSD2 (HSD11B2) (probe: 204130_at), CYP27A1 (probe: 203979_at) and the combination of both genes was analyzed with the KM plotter database (https://kmplot.com/) with recurrence free survival (RFS) as endpoint. Auto select best cut-off was chosen in the analysis. A total of 3951 breast cancer samples (all breast cancer) or 1161 patients (ER-negative breast cancer) from 50 individual datasets were split in high and low groups according to the cutoff value. The hazard ratio with 95% confidence intervals and log rank *P* value was calculated and significance was set at *P* < 0.05.

### Immunoblotting

For electrophoresis, cell lysate samples were denatured with NuPAGE LDS sample buffer and reducing agent (Thermo Fisher Scientific, Waltham, MA, USA) at 70°C for 10 min and run on 10% Bis-Tris precast gels (Thermo Fisher Scientific, Waltham, MA, USA) using NuPAGE MOPS SDS running buffer at 150 V for 1h20 min. For immunostaining and after gel electrophoresis, the proteins were transferred onto a PVDF membrane *via* semi-dry Western blotting at 25V for 2 h, blocked for 1 h at room temperature with 5 % BSA in TBS-T (10 mM Tris-HCl (pH 8.0), 150 mM NaCl, 1 % Tween 20) and stained with a monoclonal antibody against CYP27A1 (ab126785, Abcam, Cambridge, UK) diluted 1:1000, followed by the secondary anti-rabbit HRP-conjugated antibody (W401B, Promega, Madison, Wi, USA), diluted 1:10.000 in 5% BSA TBS-T. After each incubation, the membrane was washed 3 times for 5 min with TBS-T, and finally developed with Clarity Western ECL substrate (Bio-Rad, Hercules, Ca, USA) for 5 min.

### Metabolism of sterols in cells

Cells were plated into six-well plates (1 x 10^5^ cells per well for MDA-231 and HepG2 or 1.5×10^5^ cells per well for MCF7 and MDA-468) in the appropriate complete medium. One day after seeding, this medium was replaced with complete medium and cells were treated with 1 µM of either [^14^C]-5,6β-EC, [^14^C]-5,6α-EC with or without 2.5 µM tamoxifen., [^14^C]-CT (26 mCi/mmol) or [^14^C]-OCDO (26 mCi/mmol) for 24, 48 or 72 h. After incubation, media were collected and cells were washed, trypsined and counted. Lipids were extracted from both cells and media with a chloroform-methanol mixture as described previously (42). Radioactivity from organic and aqueous layers were measured using a β-counter. Lipids were then separated by TLC using as the eluent ethyl acetate for [^14^C]-5,6β-EC, [^14^C]-5,6α-EC, [^14^C]-CT and [^14^C]-OCDO or hexane/ethyl acetate for [^14^C]-cholesterol. The radioactive sterols were revealed by autoradiography. For quantification, silica zones at the expected Rf values corresponding to authentic chemical standards were scraped and radioactivity was measured using a β-counter, as previously described (15).

### Pharmacological inhibition of CYP27A1

Cells were plated into six-well plates (1×10^5^ cells per well for MDA-231 and HepG2 in the appropriate complete medium. One day after seeding, this medium was replaced with complete medium and cells were treated with 1 µM [^14^C]-CT (26 mCi/mmol) alone or in the presence of 10 µM bicalutamide. Metabolic studies were performed as described above.

### GC-MS analysis

MCF7 cells were plated in 100mm tissue culture plates at a density of 0.5×10^6^ cells per plate. Cells were allowed to adhere for 24 h and were treated by 1 µM of 5,6α-EC, 5,6β-EC, 27H-5,6EC (83% isomer α, 17% isomer β) with or without 2.5 µM tamoxifen, CT or 27H-CT for 72 h. Cell pellets were dried in a SpeedVac Savant AES 1999, DNA 130-230 (Thermo Scientific, Karlsruhe, Germany) and dried cells were weighed. 10 uL epicoprostanol (100 µg/ml dissolved in cyclohexane; Medical Isotopes Pelham, U.S.A.) as internal standard was added. 9 ml medium was reduced in a heating block under a gentle stream of nitrogen at 60°C to a final volume of about 0.5 ml. Prior to work-up 10 ul epicoprostanol (100 ug/ml dissolved in cyclohexane;) was added as internal standard. After alkaline hydrolysis (addition of 1 ml of a 90% ethanolic NaOH) all free total sterols and oxysterols were extracted by addition of 3 ml cyclohexane (2 times) and silylation of hydroxyl groups was performed at 100 °C after addition of a silylation reagent (Pyridne: Hexamethyldisilazane: Chlortrimethylsilane; 9:3:1). The TMSi-ethers were evaporated under a gentle stream of nitrogen at 100°C and the residue was dissolved in 30 µl n-decane. The same procedure was performed for all synthesized purified compounds for structural identification. First, structural identification of the synthesized standards was performed using gas chromatography - mass spectrometry in the scan mode (m/z: 150–700). Semi-Quantification of the above described isolated cell and medium TMSi-ethers was performed by gas chromatography-mass spectrometry in the selected ion monitoring mode using the characteristic ions for the individual compounds as listed in Table 1. We used the ratios between the areas of corresponding sterol/oxysterol and the internal standard epicoprostanol. The silylated sterols and oxysterols were separated on a DB-XLB (30 m length x 0.25 mm internal diameter, 0.25 µm film) column (Agilent Technologies, Waldbronn, Germany) using the 6890N Network GC system (Agilent Technologies, Waldbronn, Germany). Scans and seleced ion monitoring were performed on a 5973 Network MSD (Agilent Technologies, Waldbronn, Germany). The mass spectrometer was operated in the electron impact ionization mode. Gas chromatographic conditions were as follows: 2 µl sample was injected in the splitless mode (inlet was kept at 280°C with the helium flow at 1.1 ml/min)-The oven was first kept at 150°C for 3 min, ramped at 30°C/min to 290°C and held for 27.5 min. (43, 44)

**Table 1:** Characteristic masses and retention times of the individual trimethylsilylated sterols or oxysterols. The selected ions for quantification are marked in bold. Other are additional qualifier ions to prove the authenticity of the compound.

| Compound | Mass | Retention time (min) |
| --- | --- | --- |
| Epicoprostanol (ISTD) | **370** | 14.43 |
| 5,6β-EC | **384**, 474 | 18.2 |
| 5,6α-EC | **384**, 474 | 18.91 |
| CT | **456**, 546 | 20.3 |
| OCDO | 472, **562** | 19.0 |
| 27H-5,6β-EC | 472, 548, **562** | 28.2 |
| 27H-5,6α-EC | 472, 548, **562** | 29.5 |
| 27H-CT | 491, **544**, 634 | 31.2 |
| 27H-OCDO | 560, **650** | 31.5 |
| 25H-5,6β-EC | 472, 548, **562** | 25.3 |
| 25H-5,6α EC | 472, 548, **562** | 26.5 |
| 25H-CT | 491, **544**, 634 | 29.0 |
| 25H-OCDO | 560, **650** | 29.5 |

### Cell viability assays

MCF7 cells were seeded at 1.5 x 10^5^ cells per well in 6-well plates. After 24h, cells were treated with 1 µM of 27HC, OCDO or 27H-OCDO alone or in combination for 48 h. Cells were washed with PBS, trypsinized, resuspended in a Trypan blue solution (0.25% (w/v) in PBS) and counted in a Malassez cell under a light microscope.

### Determination of OCDO synthase activity and inhibition in cell lysates

OCDO synthase activity was measured in HEK-293T cell lysates expressing recombinant HSD2. About 5 x 10^6^ HEK-293T cells were transfected with 5 µg of the plasmid pCMV6-XL5 HSD2 (SC122552, OriGene Technologies, Inc., Rockville, MD, USA) using the Neon Transfection System (Thermo Fisher Scientific, Waltham, MA, USA) with 2 pulses at 1300 V for 20 ms. 24 h after transfection, the media was replaced by DMEM without phenol red supplemented with 10 % dextran-coated charcoal-stripped FBS. Cells were harvested 48 h after transfection, washed with PBS and stored at −80°C. For cell lysates, cell pellets were resuspended in 50 mM HEPES pH 7.4, 200 mM NaCl, 20 % glycerol, 25 mM sucrose, 5 mM DTT, 1 mM MgCl_2_, 1 mM CaCl_2_ and 1 % protease inhibitor, lysed by 4 freeze-thaw cycles in N_2_, and centrifuged for 10 min at 9000 x g. HSD2 activity was measured in a final volume of 200 µl containing 5, 10 or 20 nCi of [4-^14^C]-Cholestane-3β,5α,6β-triol (CT) to final concentrations of 0.5, 1, 1.5 or 2 µM, 0.5 mM NAD^+^, 2 % DMSO and 27H-CT at indicated concentrations. The reaction was started by adding 20 µg of cell lysate or [4-^14^C]-CT to the reaction mixture and incubated for 10 min at 37°C and stopped by adding 500 µl of ice-cold methanol. Lipids were extracted with a choloroform-methanol mixture as described before (1). The organic phase was washed with 1 ml of H_2_O to remove glycerol. The conversion rates were determined by thin-layer chromatography (TLC) using ethyl acetate as eluent. For quantification, the corresponding silica zones were scraped and radioactivity was measured using a β-counter, as previously described (15). The inhibitory constant *K*_i_ was calculated using the curve-fitting program GraphPad Prism 9.

### Lentiviral sh-RNA transduction

The plasmids pGFP-C-shLenti encoding shCYP27A1(TL313602) were from OriGene (OriGene Technologies GmbH, Herford, Ge). HEK293FT cells were seeded on T75 flasks. 24 hrs later 80% confluence HEK293FT cells were cotransfected with 10 µg of the pGFP-C-shLenti transfer plasmid, 10 µg of the p8.91 packaging plasmid and 5 µg of the pVSVg envelope plasmid using the calcium phosphate method. 6 h after transfection, the medium was replaced by 12 ml of OptiMEM medium (Life Technologies, Thermo Fisher Scientific, Waltham, MA, USA). Twenty-6 h after transfection, the conditioned medium was collected, cleared by centrifugation and filtered through 0,45 µm-pore-size PVDF filters. The lentiviral pelllet was resuspended in PBS and stored at −80°C. The cell lines MDA-231-TL313602 stably expressing shCYP27A1 (29 mer shRNA constructs against hCYP27A1 in lentiviral GFP (cat# TL313602, Origene, Rockville, MD, USA): A) TL313602A: 5’CAGGTGTCTGGCTACCTGCACTTCTTACT3’; B) TL313602C: 5’AACCAGGTGTCGGACATGGCTCAACTCTT3’; C) TL313602D: 5’GGCAACGGAGCTTAGAGGAGATTCCACGT3’, or shCont were generated by lentiviral infection. The concentrated lentivirus supernatant was aliquoted and kept at −80°C before use. For transduction experiments, MDA-231 cells were seeded onto 12-well plates (30,000 cells per well) and incubated for 24 h at 37°C with 5% CO_2_ in DMEM 10% FBS without antibiotics. The concentrated viral supernatant containing lentivirus that express shRNA targeting CYP27A1 (shCYP27A1-A, B and C) or non specific shRNA (sh-Control) were added into the culture medium at a multiplicity of infection (MOI) of 20 in the presence of protamine sulfate (5 µg/ml). One day after transduction, this medium was replaced with complete medium. After 72 h, puromycin was added to the medium at 1 μg/ml for stable CYP27A1 knock-down selection. For metabolic studies, MDA-231-shControl, shCYP27A1-A, sh-CYP27A1-B and sh-CYP27A1-C were plated into six-well plates (1×10^5^ cells per well) in the appropriate complete medium. One day after seeding, this medium was replaced with complete medium and cells were treated with 1 µM [4-^14^C]-OCDO (26 mCi/mmol), [4-^14^C]-CT (26 mCi/mmol) or [4-^14^C]-5,6α-EC (26 mCi/mmol) alone or in the presence of 2.5 µM tamoxifen for 48 h. Metabolic analyses were performed by TLC as described above. Production and cells transductions were fullfilled in the “Plateau de Vectorologie, at the “Pôle Technologique du CRCT” (Toulouse, France).

### Statistical analysis

Data are given as mean ± standard error of the mean (S.E.M) of three independent experiments each carried out in duplicate. Statistical analysis was carried out using the Student’s t-test for unpaired variables. *, **, *** and **** in the figures refer to statistical probabilities (P) of <0.05, <0.01, <0.001 and <0.0001, respectively. GraphPad Prism 9.0 software was used for all the statistical analyses.

## RESULTS

### CYP27A1 expression in breast cancer cell lines

Aiming to investigate the metabolic transformation by 27-hydroxylation of oncosterone and its precursors (5,6α-EC, 5,6β-EC and CT), we first evaluated the expression of CYP27A1 in normal mammary tissue, breast cancer and human breast cancer cell lines through the analyses of transcriptomic databases (GTEx and TNM plot). CYP27A1 is mainly expressed in the liver (332.1 transcript per million, n=226) but also in several others tissues including normal mammary tissue (47.47 transcript per million, n=459) (Fig. 3A). As previously reported (40), the analysis of transcriptomic database ([TNMplot](https://tnmplot.com/analysis/)) showed that CYP27A1 is expressed in breast cancer in paired tumors (n=7569), adjacent normal breast tissues (n=252) and metastasis (n=82), showing a slight decrease between normal and cancer tissues (Fig 3B). The cell lines used in our studies were chosen because they were reported to express CYP27A1 at various mRNA levels (https://www.proteinatlas.org/), and we confirm that at the protein level by western blotting (Fig 3C). The order for CYP27A1 expression was found as follow MCF7 (no expression) < MDA-231 < MDA-468 < HepG2 (Fig. 3C). The HepG2 cell line is an hepatoblastoma cell line that was used as a positive control known to metabolize oxysterols (45, 46). CH25H was found weakly expressed at the mRNA level in MDA-231 (https://www.proteinatlas.org/: tpm 1.2) while CYP46A1 was only detected at a very low level in MCF7 cells (https://www.proteinatlas.org/: tpm 0.3) (Supplementary Table 1). These data show CYP27A1 is widely expressed in human breast tumors while its expression can vary from null expression to high expression in cancers cell lines showing that the selected cell lines represent good in vitro models to study the 27-hydroxylation of sterols.

**Figure 3:**
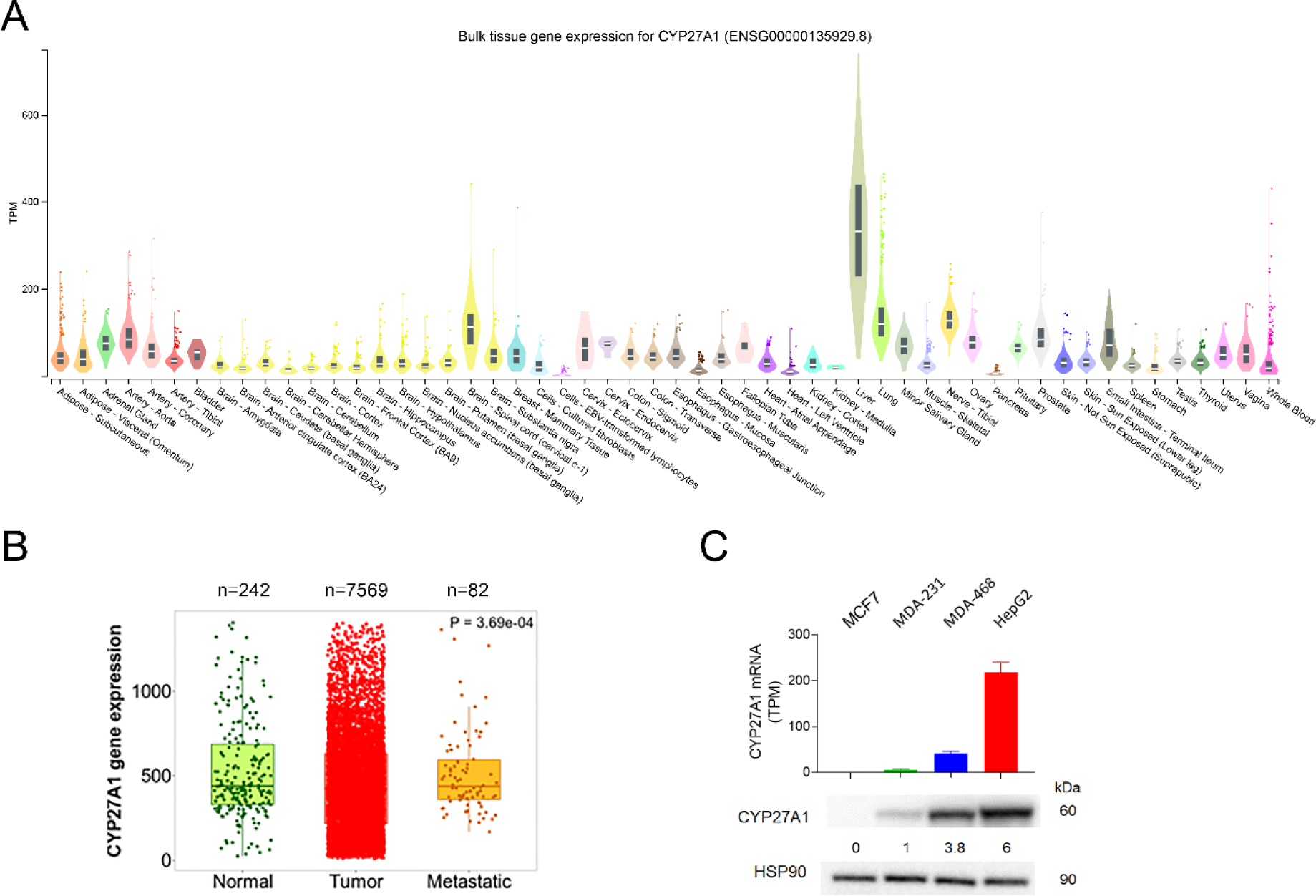
Expression of CYP27A1. A) Violin plots of bulk tissue gene expression of CYP27A1 (transcript per million; TPM) from the analysis of the transcriptomic database GTEx (GTEx Portal). B) Box plots of CYP27A1 gene expression by gene chip data in breast cancer when comparing paired normal adjacent tissue (n=242), tumor (n=7569) and metastasis (n=82) with the transcriptomic database TNM plot (https://tnmplot.com). C) CYP27A1 gene expression (transcript per million; TPM) on human breast cancer cell lines MCF7, MDA-231, MDA-468 and in the human hepatoblastoma cell line HePG2. Transcriptomic data were retrieved from the Human Protein Atlas database (https://www.proteinatlas.org/). Immunoblot of CYP27A1 protein expression in MCF7, MDA-231, MDA-468 and HepG2 cells. Western Blots are representative of three independent experiments.

### Metabolism of OCDO, CT and 5,6-EC in cancer cell lines

We first examined the metabolism of [4-^14^C]-5,6α-EC, [4-^14^C]-5,6β-EC, [4-^14^C]-CT and [4-^14^C]-OCDO over a 72-h period in the BC cell lines, MCF7, MDA-231, MDA-468 and in the hepatoblastoma cell line HepG2. Metabolic analyses were performed by thin-layer chromatography (TLC) analyses of lipidic extracts from cells and media. 25- and 27-hydroxylated synthetic compounds were used as chemical standards for the investigation of the biogenesis of side-chain hydroxylated metabolites formed from radiolabelled precursors.

#### Metabolism of OCDO

As expected in CYP27A1 positive cells (MDA-231, MDA-468 and HepG2), OCDO is converted into a compound (Rf=0.39) that comigrates with authentic 27H-OCDO, while no production was observed in MCF7 that do not express CYP27A1 (Fig. 4A). This compound was named sc-OCDO because we cannot rule out that a 25H-OCDO, which co-migrates with 27H-OCDO, could be formed in MDA-231. MDA-231 are expressing CH25H although at a low level (1.2 tpm, see supplementary Table 1). Sc-OCDO was not detected in cell extracts (data not shown) but only in culture media extracts from CYP27A1 expressing cells (Fig. 4A). In addition, we observed the appearance in the medium from MDA-468 and HepG2 of more polar OCDO metabolites than sc-OCDO (Fig. 4B). Togeteher these data showed that CYP27A1(+) cell lines secrete a metabolite of OCDO (sc-OCDO) that comigrates with authentic 27H-OCDO and which is not produced by the CYP27A1(-) MCF7 cells.

**Figure 4:**
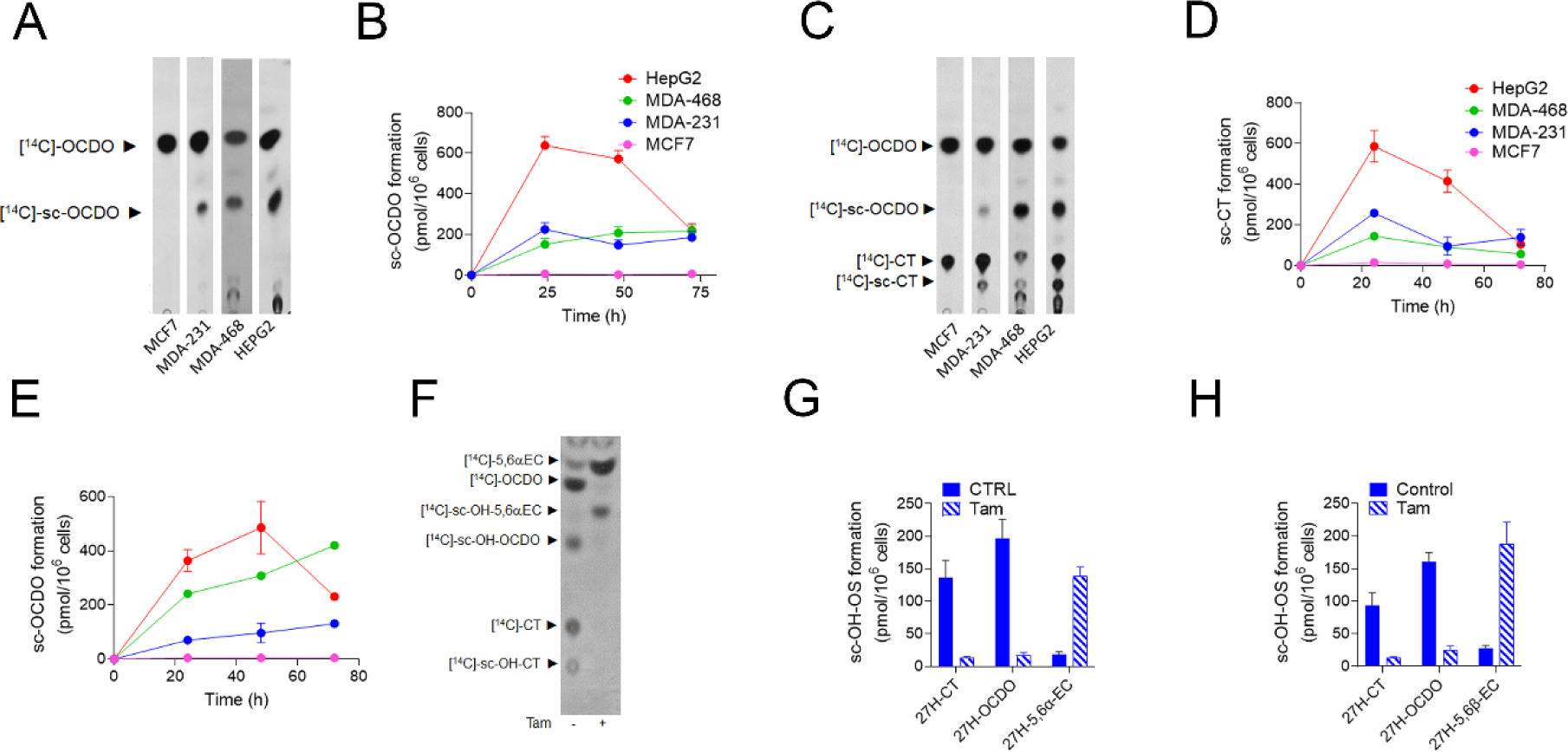
Representative TLC autoradiograms and quantitative analyses (n = 5) of lipidic extracts from cell media from cells treated with 1 μM [^14^C]-OCDO (A, B), [^14^C]-CT (C-E) for 48 h and with 1 μM [^14^C]-5,6α-EC (F, G) or 5,6β-EC (H) alone or in the presence of 2.5 µM of tamoxifen (Tam) for 72 h. Side-chain hydroxylated OCDO (sc-OCDO), sc-CT and sc-5,6-ECs co-migrates with their synthetic 25H- and 27H counterparts. For quantification, the region corresponding to sc-OCDO, sc-CT and sc-5,6-EC were recovered and counted using a β-counter. Results are the mean (±SEM) of five independent experiments.

#### Metabolism of CT

When CT was used as a substrate, we observed the production of OCDO in all tested cell lines as previously reported (5) (Fig. 4C). Both the biogenesis of sc-OCDO and sc-CT (Rf= 0.1) was visualized in the culture media from CYP27A1 expressing cells (Fig. 4C-E). Sc-CT was not oberved in the CYP27A1(-) cells MCF7. These data showed that CYP27A1 expressing cells are secreting sc-CT that is likely to correspond to 27H-CT.

#### Metabolism of 5,6-EC

Both 5,6α-EC and 5,6β-EC are hydrolyzed by ChEH to provide CT (5, 15, 36, 37) that is next oxidized into OCDO by HSD2 (5). sc-5,6α-EC or sc-5,6β-EC were not produced by CYP27A1 expressing cells in our conditions while sc-CT and sc-OCDO were present. This shows that sc-5,6α-EC or sc-5,6β-EC are rapidly consumed by ChEH (Fig. 4F-H). When the ChEH inhibitor tamoxifen (15) and 5.6-ECs were co-incubated with cells, 5,6-ECs were not further metabolized into CT and OCDO, leading to the accumulation of 5,6α-EC and 5,6β-EC and sc-5,6EC (Rf= 0.47; Fig. 4F-H), indicating that the 5,6-ECs are hydroxylated in their side chain (Fig. 4F). Together, our data show that 5,6-EC, CT and OCDO can undergo side chain hydroxylation in CYP27A1(+) cells (MDA-231, MDA-468 and HepG2) but not in the CYP27A1(-) cell line MCF7.

### Characterization and quantification of 27-hydroxylated metabolites by GC-MS

We next investigated the nature of the side chain hydroxylated metabolite produced CYP27A1-positive cell lines by GC-MS. While there was no doubt that 27H-OCDO was produced as a single sc-OCDO metabolite by MDA-468 and HepG2, MDA-231 cells expressed CH25H in addition to CYP27A1opening up the possibility that 25H-OCDO and its precursors 25H-5,6-EC and 25H-CT could be produce. Chemical standards were derivatized to their trimethylsilyl (TMS) ethers and analyzed by GC-MS as previously reported (43). We found that TMS ethers of 27H-5,6α-EC, 27H-5,6β-EC, 27H-CT and 27H-OCDO are well resolved from their 25-hydroxylated counterparts (Table 1). Fig. 5A shows the GC profile for 27H-5,6α-EC, 27H-5,6β-EC, 27H-CT and 27H-OCDO. The MS spectra corresponding to the 4 compounds are given on supplementary figures S1-S4. The GC profile of 25H-compounds and their MS spectra are given on supplementary Fig. S5-S9. Retention time and characteristic m/z data are given in Table 1. One fragment ion for each chemical standard was selected for quantification by gas chromatography– mass spectrometry selected ion monitoring method and one to two more as qualifier ions (Table 1).

**Figure 5:**
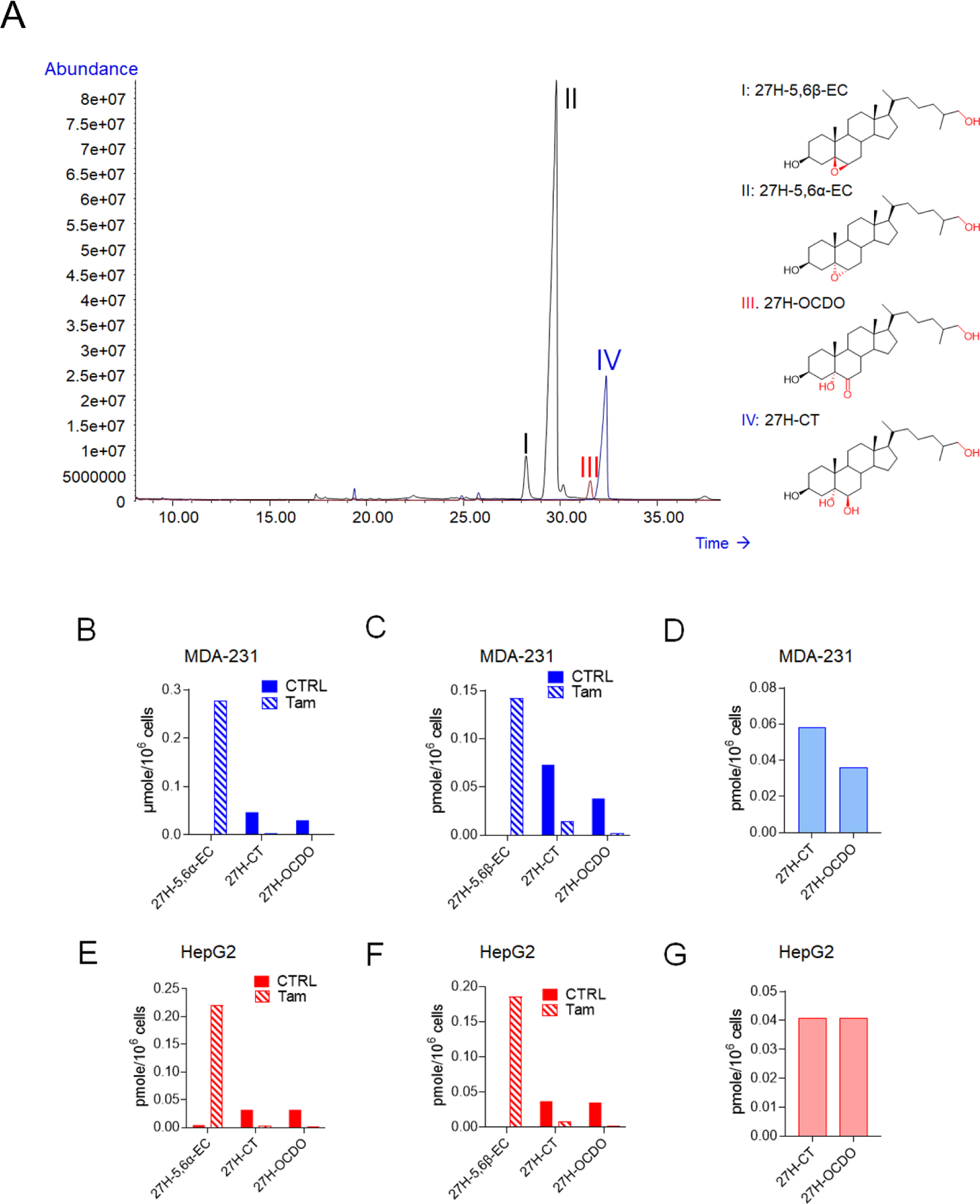
Total ion chromatogram (TIC) of the trimethylsilyl ethers (TMS) of the chemical standards including 27H-5,6β-EC (I), 27H-5,6α-EC (II), 27H-OCDO (III) and 27H-CT (IV) in full scan mode in the range of 50–750 m/z (A). GC-MS quantification of 27H-5,6α-EC, 27H-5,6β-EC, 27H-CT and 27H-OCDO in the media of MDA-231 and HepG2 cells treated for 72 h with 1 µM of 5,6α-EC (B and E), 5,6β-EC (C and F) with or without 2.5 µM tamoxifen (Tam) or with 1 µM of CT (D and G).

Then, MCF7 cells (CYP27A1(-)), MDA-231 (CYP27A1(+)) and HepG2 and MDA-468 (CYP27A1(++)) cells were exposed for 72 h to CT, 5,6α-EC and 5,6β-EC with or without the ChEH inhibitor tamoxifen. Lipid extracts from cell media were derivatized by trimethylsylilation and analyzed by GC-MS-SIM as described in Materials and Methods. As expected, 27H metabolites were not found in the CYP27A1(-) cell line MCF7, whereas both 27H-CT and 27H-OCDO were detected and quantified in the culture media of MDA-231 and HepG2 cells incubated with 5,6α-EC (Fig. 5B and 5E), 5,6β-EC (Fig. 5C and 5F) and CT (Fig. 5D and 5G). 27H-5,6α-EC or 27H-5,6β-EC were not detected in cells exposed to 5,6α-EC or 5,6β-EC respectively, except when co-incubated with tamoxifen. In that case a strong decrease in 27H-CT and 27H-OCDO levels was observed and 27H-5,6-EC appeared (Fig. 5B, 5C, 5E and 5F). These data are consistent with TLC analyses evidencing that ChEH inhibition is required to accumulate 27H-5,6α-EC and 27H-5,6β-EC. It is worthnoting that 25-hydroxylated metabolites were not detected in the cell lines used in this study. These results are in good accordance with the analysis of transcriptomic database (https://www.proteinatlas.org/) showing that MCF7, MDA-468 and HepG2 cells do not express the cholesterol-25-hydroxylase (CH25H), while MDA-231 expressed weakly CH25H (1.2 tpm) (supplementary table T1). Altogether, these experiments give the first evidence of the existence of 27H-5,6α-EC, 27H-5,6β-EC, 27H-CT and 27H-OCDO as newly identified metabolites produced in CYP27A1 expressing cells.

### 27H-CT is oxidized by HSD2 into 27H-OCDO

We showed by TLC analyses that 27H-OCDO arises from direct 27-hydroxylation of OCDO (Fig.4A and 4B). We then explored whether 27H-OCDO can also be produced by the oxidation of 27H-CT catalyzed by HSD2. We found that 27H-CT inhibited in a dose-dependent manner the conversion of [4-^14^C]-CT into [4-^14^C]-OCDO with an IC_50_ of 2.1 µM (Fig. 6A-B). Lineweaver-Burk analysis show that the Vmax is unchanged with increasing concentrations of the inhibitor (27H-CT), which is the characteristic of a competitive inhibitor. This is confirmed through a Dixon analysis (Ki=2.3 µM) (Fig. 6C-D). Therefore, 27H-CT compete with CT on the catalytic site of HSD2 involved in OCDO biosynthesis. We then evaluated by GC-MS the metabolism of 27H-CT on MDA-231 and HepG2 cells. For this purpose, cells were treated for 72 h with 27H-CT and lipidic extracts from cell media were analyzed by GC-MS-SIM as described in Materials and Methods. We observed that CYP27A1-positive cell lines metabolized 27H-CT into 27H-OCDO (Fig. 6E-F). Consequently, besides direct 27-hydroxylation, 27H-OCDO can also be produced from the oxidation of 27H-CT catalyzed by HSD2.

**Figure 6:**
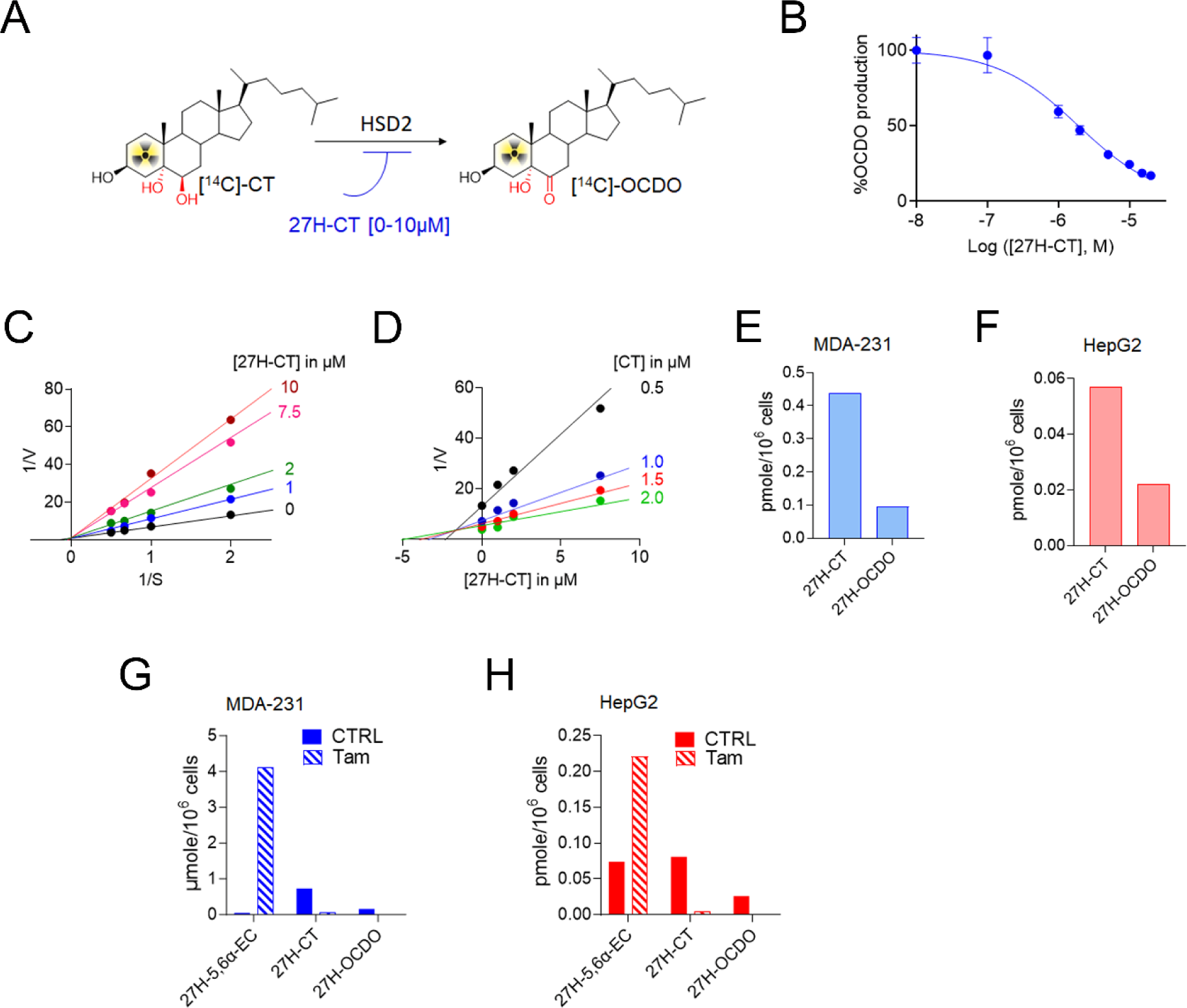
A) Scheme describing the inhibition of the OCDO synthase activity of HSD2. The OCDO synthase activity was assayed by measuring the conversion of [^14^C]-CT to [^14^C]-OCDO alone or with increasing concentrations of 27H-CT ranging from 0.1 to 20 µM. Analyses were performed by thin-layer chromatography (TLC) and quantified as described in Materials and Methods. (B) Quantification of the dose-dependent inhibition of OCDO synthase activity measured by TLC with increasing concentrations of 27H-CT and expressed as the percentage of OCDO production relative to control. The modality of the inhibition of the OCDO synthase activity inhibition by 27H-CT was done using 0, 0.1, 1, 2., 5, 10, 15 and 20 μM 27H-CT with HEK293T-HSD2 cells lysate. C) double reciprocal plots of 27OH-CT versus [^14^C]-CT. (D) dixon representation of the inhibition by 27H-CT of the OCDO synthase activity. GC-MS quantification of 27H-CT and 27H-OCDO in the media of MDA-231 (E) and HepG2 cells (F) treated for 72 h with 1 µM of 27H-CT. GC-MS quantification of 27H-5,6α-EC, 27H-5,6β-EC, 27H-CT and 27H-OCDO in the media of MDA-231 (G) and HepG2 cells (H) treated for 72 h with 1 µM of 27H-5,6-ECs (27H-5,6α-EC (83%) and 27H-5,6β-EC (17%)) with or without 2.5 µM tamoxifen (Tam).

### Metabolism of 27H-5,6-EC and 27H-CT

Metabolic studies performed by TLC and GC-MS analyses demonstrate that 27H-5,6α-EC and 27H-5,6β-EC arise from direct 27-hydroxylation of 5,6α and 5,6β-EC respectively (Fig. 4F-4H and 5). We next tested whether or not 27H-CT can also be produced by ChEH by hydrolysis of 27H-5,6-EC. MDA-231 and HepG2 cells, were treated for 72 h with 27H-5,6-EC with or without the ChEH inhibitor tamoxifen (15). Lipidic extracts from cell media were analyzed by GC-MS-SIM as described in Materials and Methods. We observed that CYP27A1-positive cell lines metabolized 27H-5,6-ECs into 27H-CT in MDA-231 (Fig. 6G) and HepG2 cells (Fig. 6H). 27H-CT is then oxidized into 27H-OCDO (Fig. 6G-H). The inhibition of ChEH by tamoxifen blocked the formation of both 27H-CT and 27H-OCDO leading to the accumulation of 27H-5,6α-EC and 27H-5,6β-EC. These data show that 27H-5,6-ECs are substrates of ChEH to produce 27H-CT.

### CYP27A1 drives the 27-hydroxylation of OCDO and its precursors

CYP27A1 is well described as the enzyme involved in 27-hydroxylation of cholesterol (47), of cholesterol precursors (48) and of the B-ring oxysterol 7-Ketocholesterol (39, 49). Consequently, we have postulated that CYP27A1 could catalyzed the biogenesis of 27H-5,6-EC, 27H-CT and 27H-OCDO from 5,6-EC, CT and OCDO respectively. The research group of Pikuleva has identified bicalutamide, as a potent inhibitors of CYP27A1 for 27HC biosynthesis (50, 51). Therefore, to confirm this hypothesis, we evaluated the impact of these compounds on the 27-hydroxylation of CT. For this purpose, MDA-231, MDA-468 and HepG2 cells were treated with [^14^C]-CT alone or in the presence of 10 µM bicalutamide. Lipidic extracts from cell media was then analyzed by TLC at the indicated time. Bicalutamide significantly reduced by approximately 60% and 40% the production of these 27-hydroxylated metabolites in MDA-231 (Fig. 7A-B) and HepG2 cells (Fig. 7C-D) respectively. This data show that the pharmacological inhibition of CYP27A1 blocked the production of 27H-CT and -OCDO.

**Figure 7:**
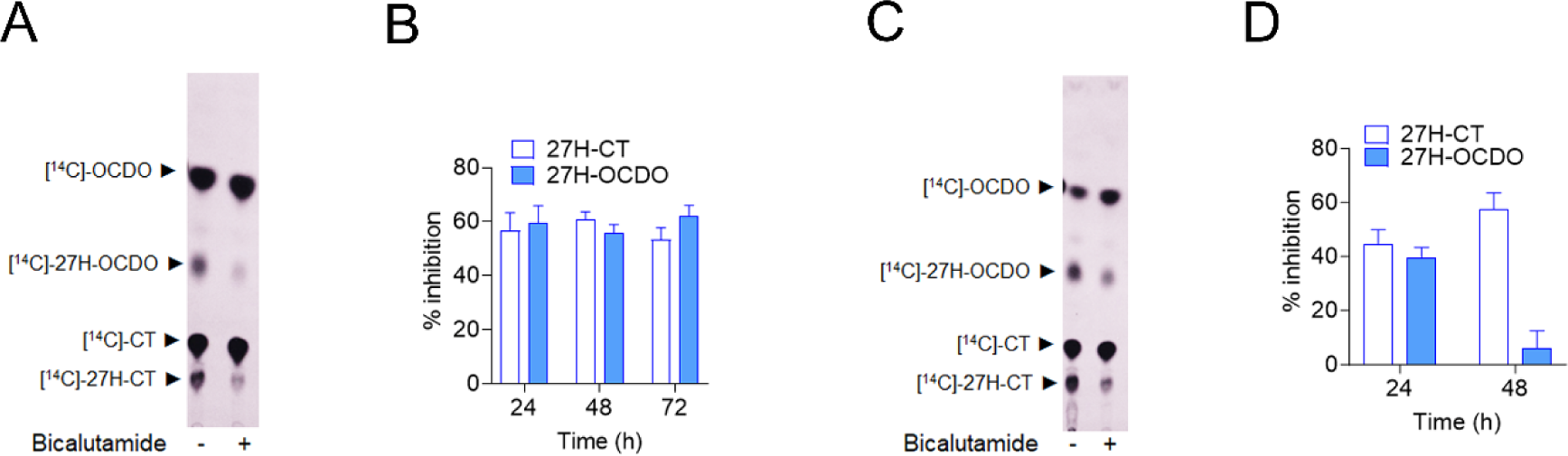
Representative TLC autoradiograms (n = 5) of lipidic extracts from cell media of MDA-231 (A) and HepG2 cells (C) treated with 1 μM [^14^C]-CT with or without 10 µM bicalutamide for 48 h. Quantitative analysis of the 27H-OCDO and 27H-CT extracted from cell media of MDA-231 (B) and HepG2 (D) and expressed at the percentage of inhibition relative to control. The region corresponding to 27H-OCDO and 27H-CT were recovered and counted using a β-counter. Results are the mean (±SEM) of five independent experiments.

To get more insight into the role of CYP27A1 in the biogenesis of 27H-5,6-EC, 27H-CT and 27H-OCDO, MDA-231 cells were transduced with three different lentivirus-shRNA against CYP27A1 (shCYP27A1-A, B and C) or with lentiviral vector carrying control non-specific sh-RNA (shControl). CYP27A1 protein level was strongly reduced in the MDA-231 cells upon treatment with the CYP27A1-targeting lentivirus (MDA-231-shCYP27A1-A, B and C) compared to MDA-231-shControl as determined by immunoblot analysis (Fig. 8A). Metabolic studies showed that CYP27A1 knock-down abolished the biogenesis of 27H-OCDO, 27H-CT and 27H-5,6α-EC on MDA-231-shCYP27A1 cells incubated with [^14^C]-OCDO (Fig. 8B-C), [^14^C]-CT (Fig. 8D-E) and [^14^C]-5,6α-EC with or without tamoxifen (Fig. 8F-G). Altogether, our data demonstrate that CYP27A1 catalyzes the 27-hydroxylation of 5,6-EC, CT and OCDO in CYP27A1 expressing cells. These data established that the genetic invalidation of CYP27A1 blocked the 27-hydroxylation of OCDO and its precusors.

**Figure 8:**
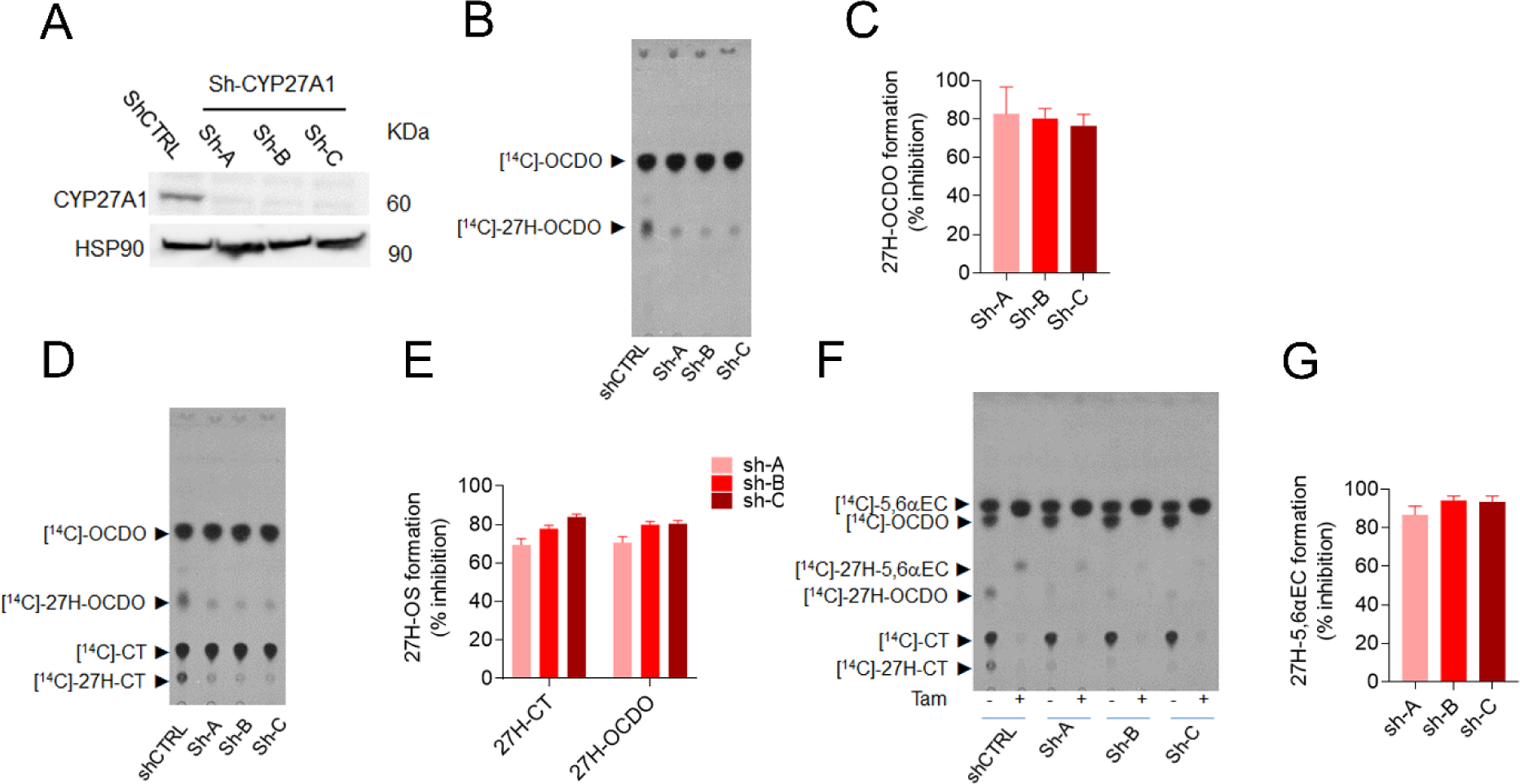
Immunoblot analysis of CYP27A1 expression in shControl (shCTRL) and shCYP27A1-MDA-231 cells (shA, shB and shC) (A) Representative TLC autoradiograms (n = 5) of lipidic extracts from cell media of shCTRL and shCYP27A1-MDA-231 cells (shA, shB and shC) treated with 1 μM [^14^C]-OCDO (B), [^14^C]-CT (D) and [^14^C]5,6α-EC in the absence or presence of 2.5 μM tamoxifen (F) for 48 h. Quantitative analysis of 27H-OCDO, 27H-CT and 27H-5,6α-EC levels expressed as the percentage of inhibition relative to shCtrl-231. The region corresponding to these metabolites were recovered and counted using a β-counter. Results are the mean (±SEM) of five independent experiments (C, E and G).

Together, these data give pharmacological and genetic evidences that CYP27A1 catalyzes the 27-hydroxylation of OCDO and its precursor.

### Comparison of the impact of OCDO and 27H-OCDO on breast cancer cells proliferation

We next investigated the impact of 27H-OCDO on BC cells proliferation alone or in combination with OCDO or 27HC. As expected OCDO and 27HC stimulated MCF7 (ERα(+)) cells proliferation as reported (5, 22, 23) while 27H-OCDO inhibited their proliferation (Fig. 9A and 9C). In addition, 27H-OCDO significantly inhibited BC cells proliferation induced by OCDO or 27HC (Fig. 9A and 9C). Moreover, we found that OCDO, but not 27HC, stimulated the proliferation of MDA-231 (TN) cells (Fig. 9B). Similarly to what was found on MCF7 (ER(+)) cells, 27H-OCDO blocked the effect of OCDO (Fig. 9B). We previously described that high levels of HSD2 (*HSD11B2*) mRNA were significantly associated with a poor prognosis (decreased recurrence free survival rate) in all BC patients (Fig. 9D). Interestingly, a Kaplan-Meier analysis of high versus low expression of CYP27A1 increases the recurrence free survival in all BC (Fig. 9E) and this effect is also observed considering the combination HSD11B2/CYP27A1 (Fig. 9F). These effects are even more pronounced considering TN BC only (Fig. 9G-I).

**Figure 9:**
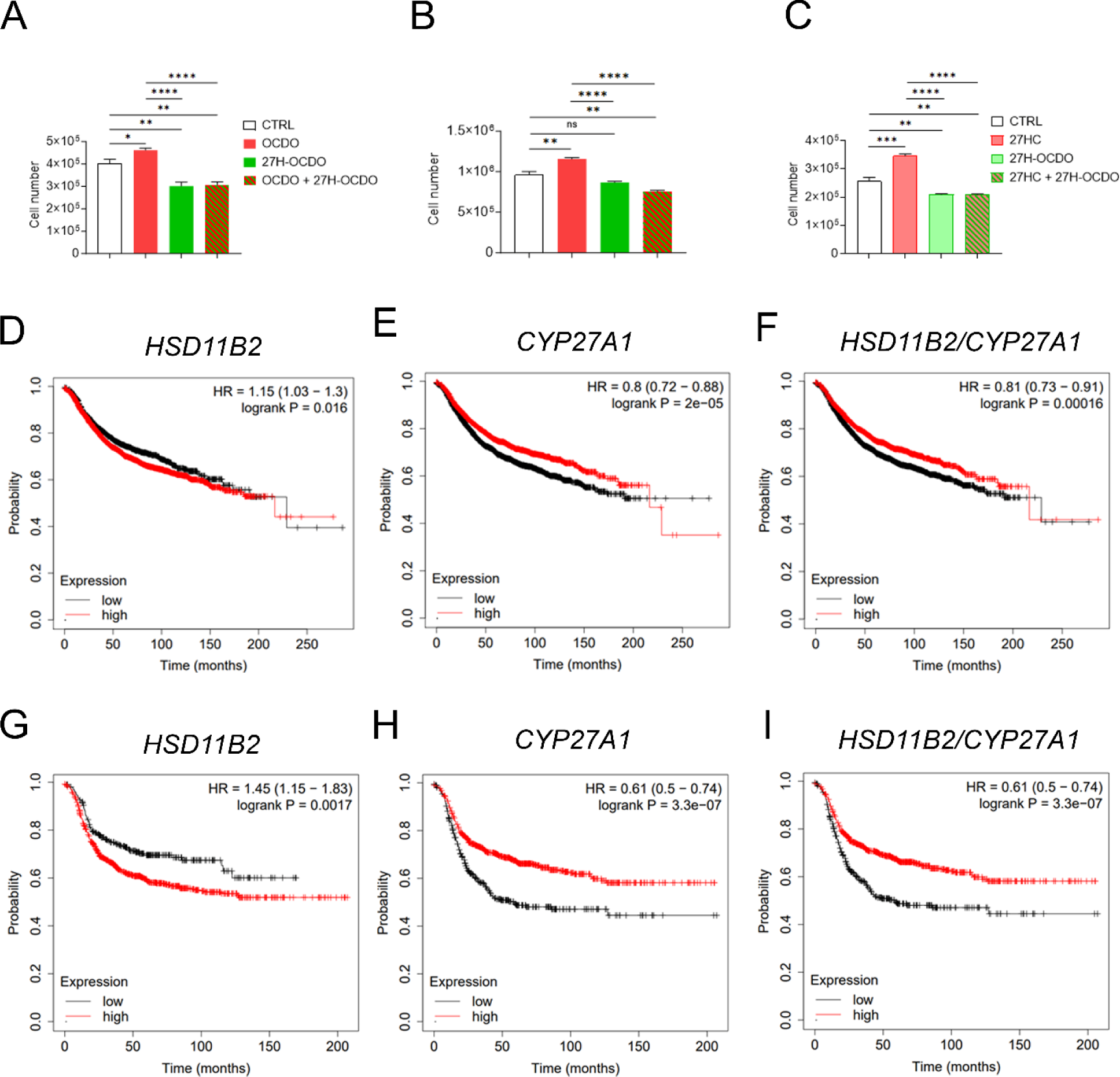
MCF7 and MDA-231 cells were treated or not for 48 h with OCDO (1 µM), 27HC (1 µM) or 27OH-OCDO (1 µM) alone or in combination. Cells were harvested by trypsinization and counted on a counting chamber Mallassez as described in Materials and Methods. Cell number are expressed as the percentage relative to control (vehicle) (A-C). Kaplan–Meier representation of patient recurrence free survival taking into account the expression (median cut-off) of the HSD11B2 (D), CYP27A1 (E) or HSD11B2 and CYP27A1 (F) genes using the Kmplot database on 50 individual datasets (4,934 samples) on all BC. Kaplan–Meier representation of ER-negative breast cancer patient recurrence free survival taking into account the expression (median cut-off) of the HSD11B2 (G), CYP27A1 (H) or HSD11B2 and CYP27A1 (I) genes using the Kmplot database on 50 individual datasets (1161 samples). Survival curves are based on Kaplan–Meier estimates and log-rank P values were calculated for differences in survival. Cox regression analysis was used to calculate hazard ratio (HR).

These data established that 27-hydroxylation of OCDO converts the tumor promoting property of OCDO into a compound with antiproliferative properties. In addition, 27H-OCDO inhibits the proliferative activities of both 27HC and 27H-OCDO. These effects are consistant with the protective effect found when CYP27A1 and HSD2 are overexpressed in all BC patients and TN BC.

## DISCUSSION

We report herein the chemical synthesis of 25- and 27H-5,6-EC, -CT and -OCDO. Using these compounds as standards, we were able to show that 27H-5,6-EC, 27H-CT and 27H-OCDO are produced in CYP27A1 expressing cells. Their identity was confirmed using chromatographic, pharmacological and genetic approaches. [^14^C]-labelled-intermediates in the synthesis of OCDO gave compounds that co-migrates with the 25H- and 27H-standards in TLC in cells expressing CYP27A1 such as MDA-231, MDA-468 and HepG2 cells but not in MCF7 cells that do not express CYP27A1. Pulse done with unlabelled precursors and analyses by GC-MS-SIM of cell media extracts established that only 27H-5,6-ECs, 27H-CT and 27-OCDO compounds were produced, ruling out the production of 25H-metabolites by these cells. We observed that these metabolites are mainly secreted in cell culture media and weakly or not detectable in cells showing that they are diffusible metabolites. 27H-5,6-ECs were detected only when ChEH activity was pharmacologically inhibited by drugs. Our data demonstrate for the first time that 27-hydroxylated 5,6-ECs, CT and OCDO exist as human metabolites. We also elucidate the metabolic pathways involved in the biogenesis of these metabolites. Indeed, we showed that 27-hydroxylated 5,6-EC, CT and OCDO are produced by direct 27-hydroxylation catalyzed by CYP27A1 (Fig. 8 and 10). In addition, 27H-OCDO is formed by oxidation of 27H-CT driven by HSD2 whereas ChEH catalyzes the hydrolysis of 27H-5,6α and β-EC towards 27H-CT (Fig. 10).

**Figure 10:**
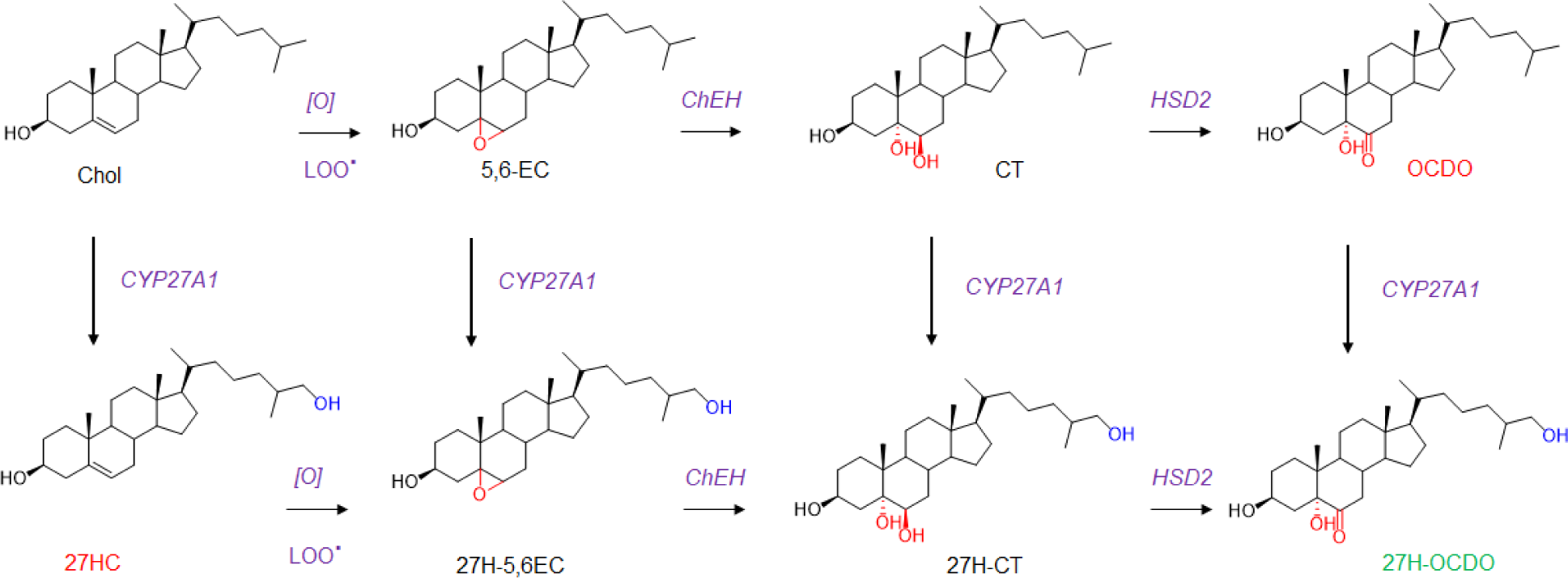
Metabolic pathways showing the biogenesis of 27H-5,6-EC, 27H-CT and 27H-OCDO. [O] and LOO∙ represent respectively oxidative stress and lipid hydroperoxides radicals involved in the epoxidation of delta-5,6 double bond of cholesterol and 27HC. 27HC and OCDO are BC tumor promoters that induce proliferation (in red) whereas 27H-OCDO is antiproliferative and inhibits proliferative properties of 27HC and OCDO (in green).

It is highly probable that 27HC undergoes epoxidation at the level of its delta-5,6 double bond to give 27H-5,6α and β-EC through lipoperoxidation or via cytochrome p450 as previously reported for cholesterol (35). Interestingly, we found that an elevated CYP27A1 expression in cells (MDA-468 and HepG2) leads to further metabolic transformations of 27H-5,6-EC, 27H-CT and 27H-OCDO into more polar metabolites. Since we showed that the pharmacological and the genetic inhibition of CYP27A1 inhibited their production, they are probably bile acid derivatives of 27H-compounds. Several studies showed that sterol and oxysterol metabolism in HepG2 cells led to lipid soluble C24 and C27 bile acids as well as their water soluble sulfate esters. It is thus highly probable that 27H-EC, 27H-CT and 27H-OCDO are converted into C24 and C27 bile acids (lipid soluble metabolites) and their sulfate ester (water soluble metabolites) as observed for 7-ketocholesterol in HepG2 cells (39, 46, 49, 52), We previously reported that OCDO is a tumor promoter on ERα(+) and TN BC inducing cancer cells proliferation via the glucocorticoids receptor (5). In addition, it was shown that OCDO level as well as the expression of its biosynthetic enzymes (ChEH and HSD2) are increased in BC compared to normal adjacent tissue and are positively associated with a poor prognostic (5). 27-HC was reported to stimulate the proliferation of ERα(+) BC cells via ER, to induce an epithelial-mesenchymal transition and to promote an immunosuppressive tumor microenvironment through LXR favoring BC metastasis spreading (22, 23, 53). However, epidemiologic studies from the EPIC-Heidelberg cohort showed that higher circulating 27HC was associated with lower risk of breast cancer in postmenopausal women (26, 54). Interestingly, we found that 27H-OCDO does not stimulate cell proliferation, but instead displays anti-proliferative properties *in vitro* and inhibits the stimulation of BC cells proliferation by OCDO and 27HC. Thus, 27H-OCDO level could better correlate with patient survival. The characterization of these new metabolites arising from 27HC and from the 5,6-EC metabolic pathway offers new opportunities to get a better understanding of the relationship between oxysterols metabolism and BC development and will provide new standards and quantitative methods that will be available for epidemiological studies. Our data established that 27HC and OCDO are not end metabolic products and showed that these pathways can cross together to give mixed side-chain and B-ring oxysterols with antiproliferative properties. Our data suggest that directing BC cell metabolism toward 27H-OCDO could switch the pro-tumor activities of both 27HC and OCDO into potential endogenous anti-tumor compounds, opening up new perspective in our understanding of BC occurrence and development. Since 27H-OCDO is the end-product in cells expressing low CYP27A1 levels but not in cells expressing high CYP27A1 levels, it will be of interest to determine the chemical structure and the biological properties of its bile acid derivatives. The accumulation of 27HC has been measured in the sera of patients suffering from Spastic paraplegia type 5 (SPG5), a genetic neurological disorder caused by recessive mutations in the gene CYP7B1 that is involved in 27HC catabolism (55). Whether 27H-5,6-EC, 27H-CT and 27H-OCDO also accumulate in SPG5 and their impact on neurodegeneration related to this genetic syndrome deserve to be studied. In addition, the relationship between CYP7B1 expression in BC and the level of these metabolites warrants investigation. 25H-metabolites were not detectable in the cell lines used in this study that do not (MCF7, MDA-468 and HepG2) or weakly (MDA-231) express CH25H. However, transcriptomic database showed that some human BC cell lines (i.e., MDA-157: tpm=2.8, MDA-436: tpm=26.9, https://www.proteinatlas.org/,) but also BC from patients, normal adjacent tissues and metastasis express CH25H ([TNMplot](https://tnmplot.com/analysis/)). Consequently, the biogenesis of 25H-OCDO and its metabolic precursors probably occurs in CH25H expressing cells. CH25H is notably highly expressed in immune cells including macrophages and leukocytes and its expression is increased by inflammation (56). It is noteworthy that CH25H have been reported for its key role in the control of inflammation and antitumor immune response. For example, in pancreatic cancer, the loss of CH25H expression is associated with downregulation of MHC-I and decreased CD8^+^ T-cell tumor infiltration (57). In addition, the impairment of cytotoxic T lymphocytes (CTLs) activity induced by trogocytosis between malignant cells and CTLs involve the downregulation of CH25H expression in various solid tumors (58). It was also reported that low expression of CH25H in leukocytes was significantly associated with lung adenocarcinoma metastasis (59). Consequently, whether 25H-OCDO is produced in BC and more particularly in tumor infiltrated immune cells, its impact on BC cells proliferation and invasiveness as well as on antitumor immune response deserve to be investigated in the future. This study provides both chemical standards and quantitative GC-MS method to quantify 25H-OCDO and its precursors in biological samples. In conclusion, this study identified new human cholesterol metabolites highlighting the importance of the 5,6-EC branch on the cholesterol pathway and in BC progression or control.

## AUTHOR CONTRIBUTION

Conceptualization, M.P., S.S.-P., and P.D.-M.; Methodology, P.D.-M. did the chemical synthesis the characterization and the purification of unlabelled compounds. P.D.-M., S.A., and R.S did the the radiolabelled chemistry; P.D.-M., L.P. and S.A. did the biochemistry, the enzymology and cell proliferation experiments. S.F. and D.L. did GC/MS experiments.; Writing, P.D.-M., S.S.P. and M.P. and C.J.V.; Reviewing: P.D.-M., S.A., D.L., S.S.-P. and M.P.; Funding Acquisition, M.P. and S.S.-P..; Supervision, M.P., S.S.-P. and P.D.-M..

## DATA AVAILABILITY

Data will be shared upon written request to Marc Poirot at CRCT through email

## ACKNOWLEDGEMENTS

We thank Dr Loïc Van den Berghe from the Pole Technologique at CRCT for the amplification of plasmids, the production of virions and for his advice for transfection experiments. Structural analyssis of compounds were done at the “Service Commun de Résonance Magnétique Nucléaire - Plateforme scientifique et technique ICT FR 2599 UPS Module de Haute Technologie and at the ICT - UPS CNRS - Service Commun de Spectrométrie de Masse from the université of Toulouse III - Paul Sabatier, Toulouse, France. This work was funded by an internal grant from the “Institut National de la Santé et de la Recherche Médicale”, the “Université de Toulouse III”, the « Agence Nationale de la Recherche » (DASYNT2, ANR-20-CE11-0005) and the Institut National du Cancer (PLBIO-2018-145). This study has been partially supported through the grant EUR CARe N°ANR-18-EURE-0003 in the framework of the Programme des Investissements d’Avenir.

