## supplementary data for "27-hydroxylation of oncosterone by CYP27A1 switchs its activity from pro-tumor to anti-tumor"

*^1^ Cancer Research Center of Toulouse (CRCT), Inserm, CNRS, University of Toulouse, Team INOV:"Cholesterol Metabolism and Therapeutic Innovations”, Toulouse, France.^2^ Equipe labellisée par la Ligue Nationale contre le Cancer.^3^ French network for Nutrition physical Acitivity And Cancer Research (NACRe network), Jouy en Josas, France. ^4^ Institute of Clinical Chemistry and Clinical Pharmacology, University Hospital Bonn, Bonn, Germany.*

Supplementary data S1

Lipid maps codes for new compounds

LMST01010580 | 27-hydroxy-OCDO

LMST01010579 | 25-hydroxy-OCDO

LMST01010581 | 25-hydroxy-cholestanetriol

LMST01010582 | 27-hydroxy-cholestanetriol

LMST01010583 | 25-hydroxy-5,6beta-epoxycholesterol

LMST01010584 | 25-hydroxy-5,6alpha-epoxycholesterol

LMST01010585 | 27-hydroxy-5,6alpha-epoxycholesterol

LMST01010586 | 27-hydroxy-5,6beta-epoxycholesterol

Supplementary Table 1

Expression in mRNA of enzymes. Data are takeen from the Human Protein Atlas (https://www.proteinatlas.org/)

| mRNA (T.P.M.) | MCF7 | MDA-MB231 | MDA-MB468 | HepG2 |
| --- | --- | --- | --- | --- |
| CYP27A1 | 0.0 | 5.1 | 39.4 | 217.7 |
| CH25H | 0.0 | 1.2 | 0.0 | 0.0 |
| CYP46A1 | 0.3 | 0.0 | 0.0 | 0.0 |

**Supplementary figure S1**. Mass spectrum of compound I corresponding to the TMS derivative of 27H-5,6ß-EC at the average retention time of 28.07 to 28.35 min.

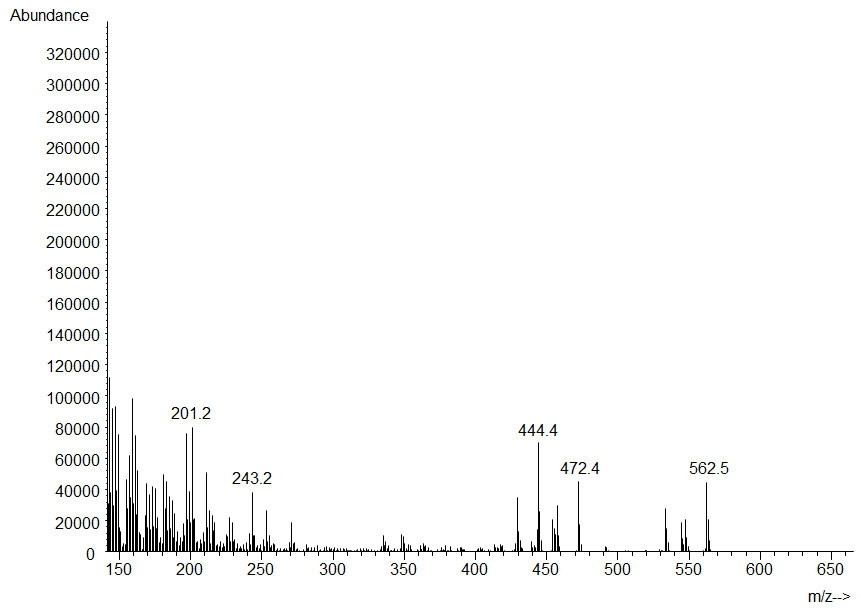

**Supplementary Figure S2:** Mass spectrum of compound II corresponding to the TMS derivative of 27H-5,6α-EC at the average retention time of 29.17 to 29.81 min.

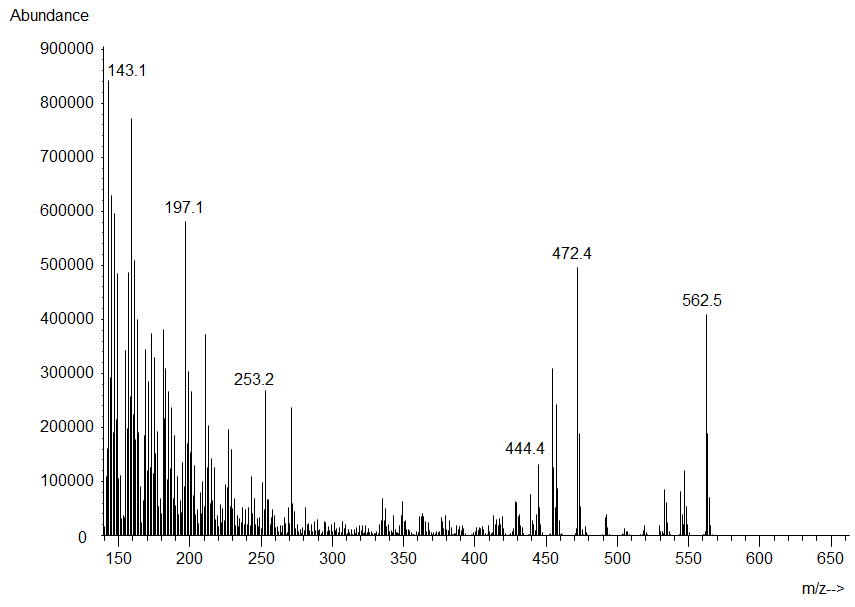

**Supplementary Figure S3:** Mass spectrum of compound III corresponding to the TMS derivative of 27H-OCDO at the average retention time of 31.27 to 31.73 min.

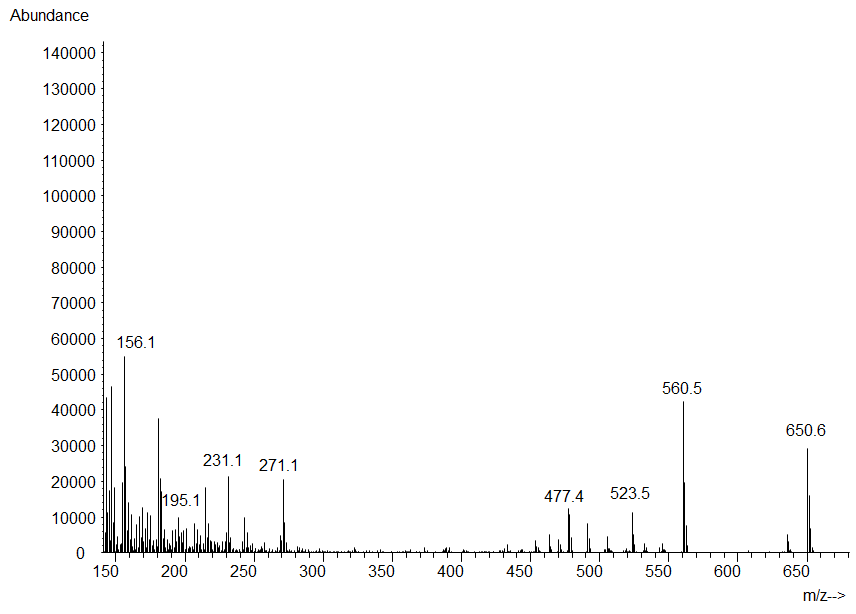

**Supplementary Figure S4:** Mass spectrum of compound IV corresponding to the TMS derivative of 27H-CT at the average retention time of 31.75 to 32.41 min.

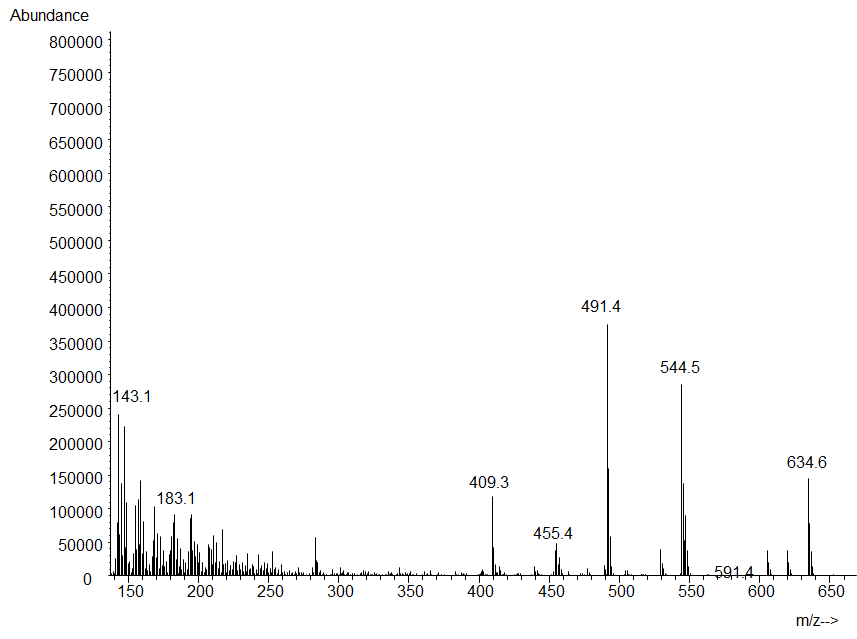

**Supplementary Figure S5**: Total ion chromatogram (TIC) of the trimethylsilyl ethers (TMS) of the chemical standards including 25H-5,6α-EC (I), 25H-5,6β-EC (II), 25H-OCDO (III) and 25H-CT (IV) in full scan mode in the range of 50–750 m/z.

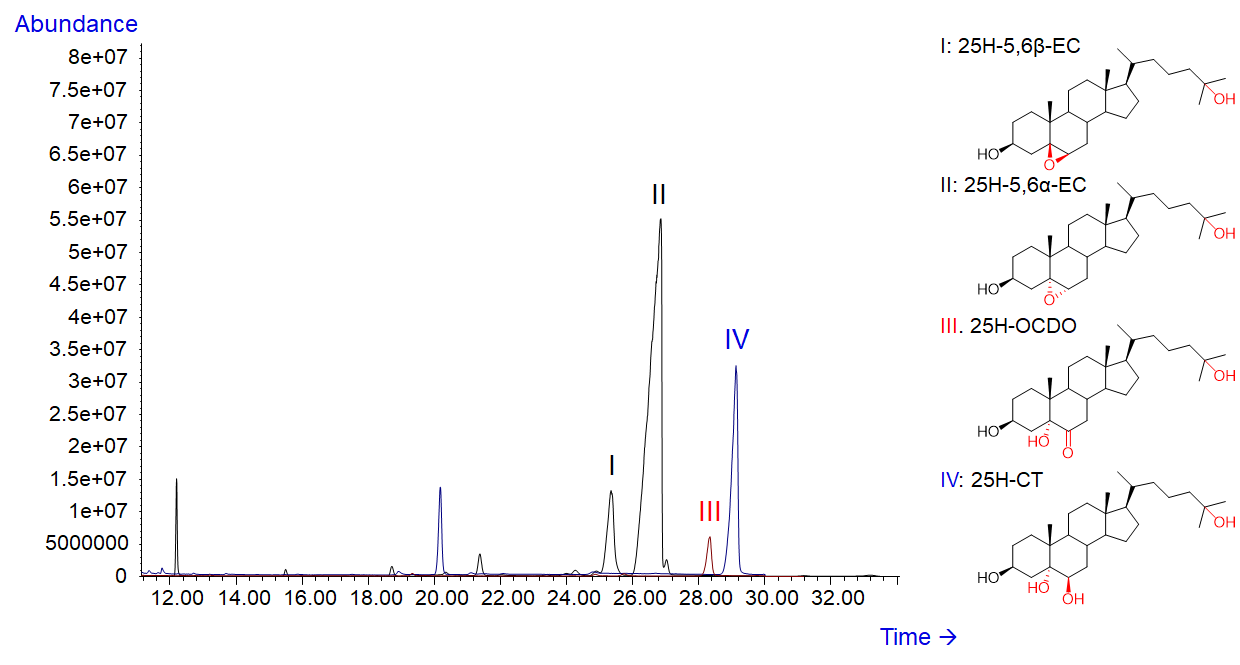

**Supplementary Figure S6:** Mass spectrum of compound I corresponding to the TMS derivative of 25H-5,6ß-EC at the average retention time of 25.11 to 28.47 min.

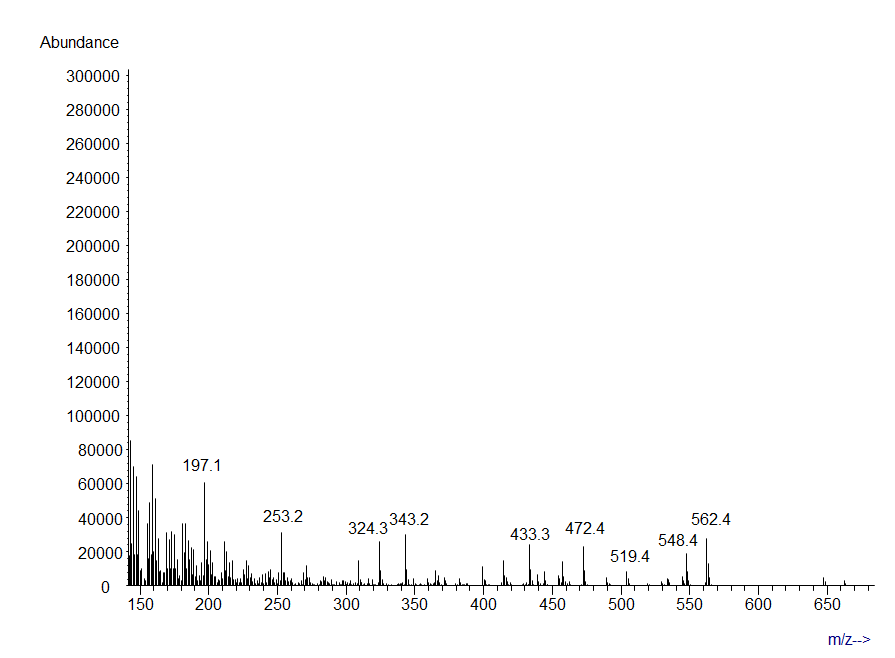

**Supplementary Figure S7:** Mass spectrum of compound II corresponding to the TMS derivative of 25H-5,6α-EC at the average retention time of 26.13 to 26.83 min.

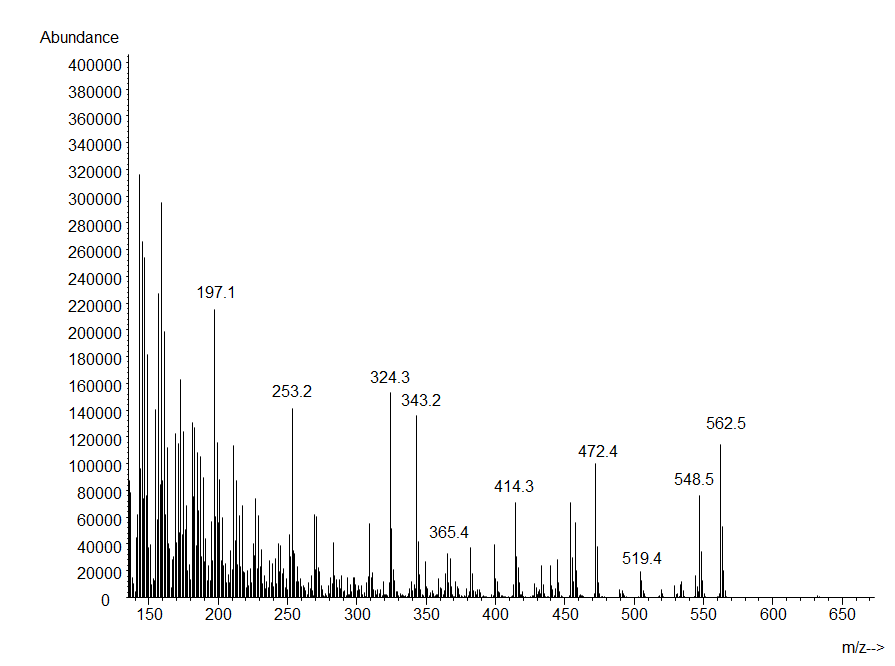

**Supplementary Figure S8:** Mass spectrum of compound III corresponding to the TMS derivative of 25H-OCDO at the average retention time of 28.06 to 28.46 min.

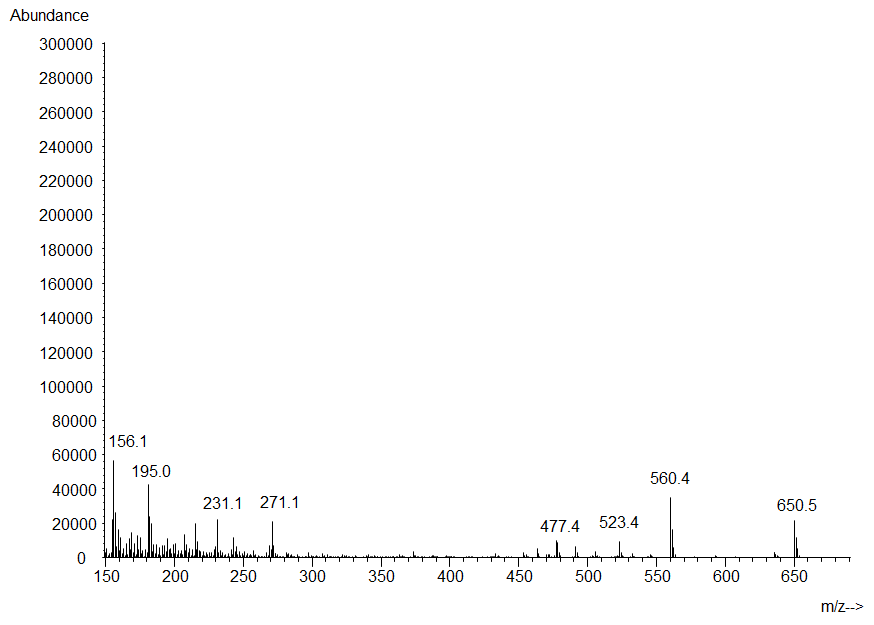

**Supplementary Figure S9:** Mass spectrum of compound IV corresponding to the TMS derivative of 25H-CT at the average retention time of 28.06 to 28.46 min.

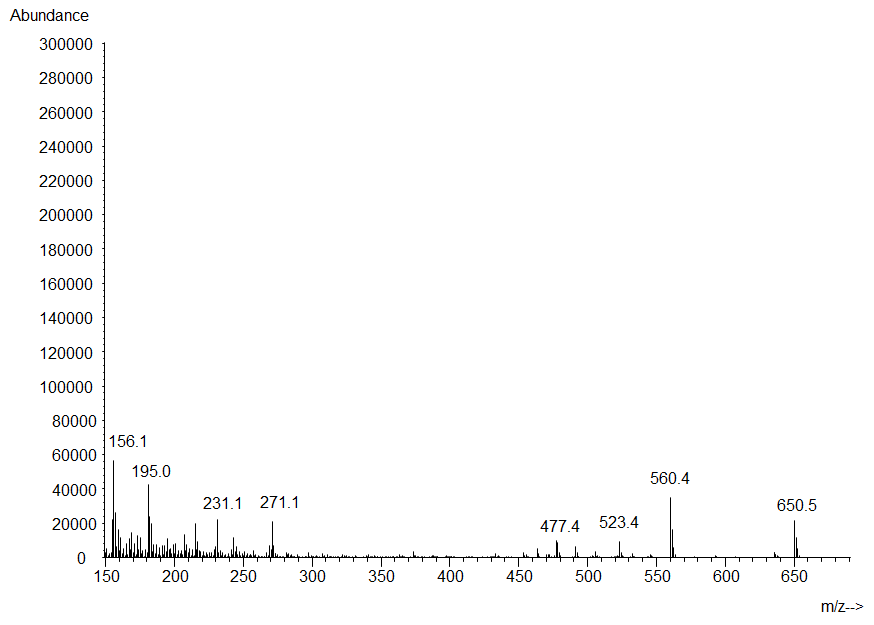
